# Beyond the blood: Tissue-resident immunity shapes SARS-CoV-2 vaccine antibody responses

**DOI:** 10.64898/2026.07.23.740375

**Authors:** Juliane Schröter, Christiaan H. van Dorp, Julia Davis-Porada, John R. Teijaro, Donna L. Farber, Andrew J. Yates

**Affiliations:** Department of Pathology and Cell Biology, Columbia University Irving Medical Center, New York City, NY, USA; Department of Microbiology and Immunology, Columbia University Irving Medical Center, New York City, NY, USA; UCLA David Geffen School of Medicine, Los Angeles, CA, USA; Department of Immunology and Microbiology, The Scripps Research Institute, La Jolla, CA, USA

**Keywords:** COVID-16, SARS-CoV-2, vaccination, adaptive immunity, humoral immunity, tissue-resident immunity, cell-mediated immunity, memory T cell, memory B cell, mechanistic inference

## Abstract

Infections and vaccinations elicit coordinated humoral and cellular adaptive immune responses that together provide protection. In addition to antibodies, pathogen-specific memory T and Bcells persist in blood and tissues, but it remains unclear how their composition and spatial distribution relate to serum antibody titers, the most common correlate of vaccine-induced protection. Understanding these relationships is essential for predicting vaccine efficacy and optimizing immunization strategies. We analyzed tissues from 58 adult human organ donors vaccinated against SARS-CoV-2, including individuals with and without prior infection. Using multivariate imputation, dimensionality reduction, and correlation, regression, and causal analyses, we identified immune signatures linking memory B cell, CD4 T cell, and CD8 T cell subsets in spleen, lung, and lung-draining lymph nodes with antibody titers and neutralizing activity. Our analyses indicate that humoral immunity is driven primarily by virus-specific B cells and CD4 T cells in lymphoid tissues rather than blood, whereas tissue-localized CD8 T cell responses, although correlated with antibody levels, develop independently. These findings demonstrate that cross-sectional immune profiling across multiple tissues recapitulates established immunological principles and reveal that serum antibody responses emerge from coordinated cellular immune responses distributed throughout the body.

## Introduction

Protective immunity induced by vaccination is commonly evaluated using circulating antibody titers, including binding antibodies (bAbs) and neutralizing antibodies (nAbs) – bAbs measure antigen recognition, whereas nAbs reflect the ability to directly inhibit pathogens. These antibody measurements are reproducible, scalable, and correlate with clinical protection across diverse infectious diseases (1, 2). Serological assays are also relatively easy to perform and highly standardized, making antibody titers widely used as practical surrogate measures of immune protection. As a consequence, these assays became the dominant tools for assessing SARS-CoV-2–specific immunity during the COVID-19 pandemic (3–7). The pandemic generated extensive serological datasets enabling quantification of immune protection (8), characterization of waning immunity (6, 10), and evaluation of vaccine performance across platforms, populations, and time since immunization (11–13). Consistent with other infectious diseases, SARS-CoV-2–specific bAb and nAb titers correlate inversely with infection risk and disease severity at the population level (8, 14).

However, serum antibody titers do not fully capture protective immunity. Breakthrough infections occur despite detectable bAbs (15), and clinical observations suggest that impaired B cell responses do not translate into proportionally worse outcomes in many settings (16). Protective immunity is therefore increasingly understood as arising from a coordinated and spatially distributed system, of which circulating antibody levels are only one measurable output. For example, antibody responses are sustained by plasma cells and memory B cells (mBcs), but their generation, maturation, and maintenance depend on interactions with CD4 T follicular helper (T_FH_) cells and regulatory T cells (T_reg_). In parallel, other CD4 T cell responses coordinate the recruitment and activity of multiple immune cell types, and CD8 T cells limit viral replication through both cytotoxic and other mechanisms.

It has become increasingly clear that the major reservoirs for T and B cell memory are maintained in tissues, including lymphoid organs and mucosal sites, rather than the blood (BLD) (17–16). Following respiratory virus infection, tissue-resident memory T cells (T_RM_) can be established in the lung (LNG) and airway mucosa, where they provide rapid local protection upon reinfection. Likewise, key components of humoral immunity – including mBcs, plasma cells, and T_FH_ cell niches – are maintained within lymphoid organs. While these tissue-resident populations are well characterized in experimental models, their contributions to protective immunity in humans remain difficult to assess because tissue sampling is rarely feasible beyond limited approaches such as lymph node (LN) aspiration.

Consequently, relatively few studies have examined SARS-CoV-2 immunity directly in tissues. For example, Ng *et al.* (20) analyzed cellular immune responses in the LNG and gut of a patient who died from severe COVID-16, revealing marked tissue-specific immune activation. Using a unique tissue resource recovered from previously healthy deceased donors, we identified T cells specific for SARS-CoV-2 epitopes in LNG and LNs up to six months after natural infection, showing correlations with circulating responses but marked tissue-specific functional heterogeneity (21). More recently, we drew on the same resource to demonstrate that vaccine-induced SARS-CoV-2-specific T cell subsets persist throughout the body, with effector responses enriched in circulation and regulatory programs enriched in tissues (22). Nevertheless it remains unclear how the phenotypic diversity and spatial distribution of the adaptive immune response to SARS-CoV-2 relates to serum-antibody measures of protection. Studying these relationships is further complicated by inter-individual variation in the timings and strains of previous infections, vaccination history, and demographics (23).

To address this gap, we analyzed data from a unique cohort of 58 deceased adult organ donors from our tissue resource, all vaccinated against COVID-19 and a subset of whom were previously infected with SARS-CoV-2 (22). This resource enabled comprehensive profiling of SARS-CoV-2-specific immune responses across seven anatomically distinct tissues – including mucosal and lymphoid sites – alongside matched peripheral BLD. For each donor, we quantified serum antibody titers and tissue frequencies of SARS-CoV-2–specific T and B cell subsets. We integrated these data using a variety of statistical approaches to identify a consensus set of immune features and demographic factors associated with inter-individual variation in serum antibody responses. We then used structural equation modeling to evaluate putative causal structures underlying these associations. These analyses support a framework in which demographics and antigen exposure combine with coordinated T and B cell responses, to a greater extent within tissues than in BLD or bone marrow (BM), to drive serum antibody responses.

## Results

### Cohort overview reveals high inter-individual variability

We analyzed data from 58 adult human organ donors vaccinated against SARS-CoV-2, obtained from up to seven distinct tissues or sites (Figure 1A, Methods). Donors were aged between 23 and 85 years, with 38 having a history of natural SARS-CoV-2 infection and 20 uninfected. Further sociodemographic data are given in Figure 1B and SI-Table S1. The dataset comprised frequencies of SARS-CoV-2 spike-reactive B and T cell subsets within tissues (Figure 1C, SI-Figure S1), measured using the Antigen-Induced Marker (AIM) assay (Methods), together with phenotypic characterization of these populations. Serum measurements included circulating Spike (S)-specific and Receptor-Binding Domain (RBD)-specific IgG antibody titers (Figure 1D, Methods), as well as neutralization titers against the ancestral Wuhan (WA1, n=53), Delta (n=48), and Omicron BA.2.12.1 (Omicron, n=49) SARS-CoV-2 variants (Figure 1E, Methods). Almost all measurements demonstrated substantial inter-individual variability. Serum-derived data were obtained for 57 of 58 donors (287 antibody measurements). Cellular response data comprised a total of 7,308 expected measurements across 58 donors and 126 cellular subpopulations, but were incomplete for many individuals due to unavailability of certain tissue samples (n=2,712), insufficient cell numbers (n=763), or left-censoring below the detection limit of the AIM assay (n=161). Consequently, each donor exhibited a unique pattern of measurements across 126 cellular subpopulations. A subset of the cellular and serological measurements from this cohort has been reported previously in (22). The present study expands that work by including additional donors, expanded cellular phenotyping, and newly generated neutralization data.

**Figure 1:**
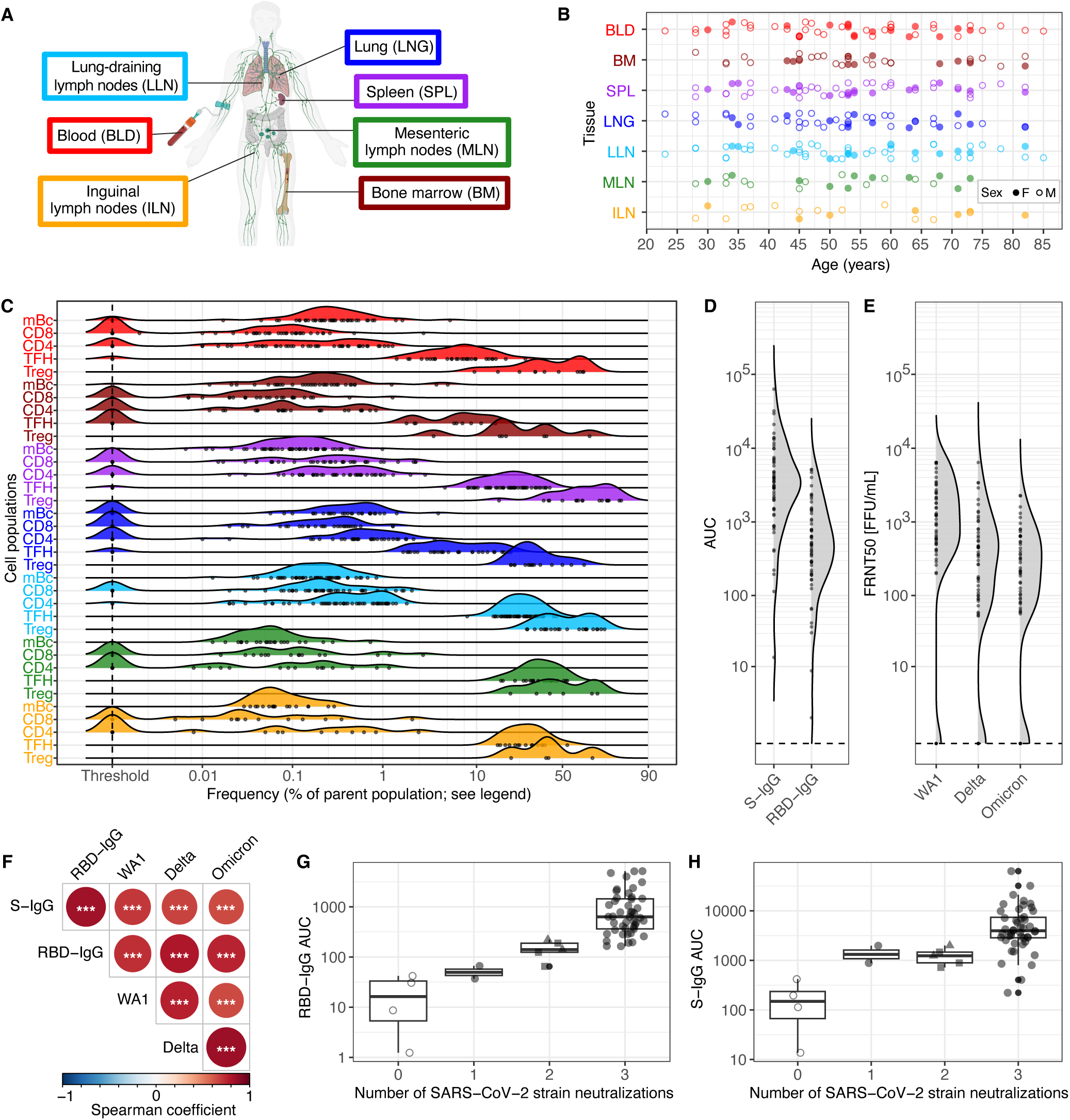
Study cohort and dataset overview: Cohort characteristics, tissue sampling, and SARS-CoV-2–specific immune readouts. (A) Schematic overview of the selected tissues and introduction of the tissue color scheme used consistently throughout the manuscript. Created with BioRender.com. (B) Tissue availability by donor age and sex. (C) Percentages of Spike-reactive (S+) cellular immune responses across tissues on a logit scale (shown for: blood (BLD; red), bone marrow (BM; dark red), spleen (SPL; purple), lung (LNG; blue), lung-associated lymph nodes (LLN; light blue), mesenteric lymph nodes (MLN; green), and inguinal lymph nodes (ILN; yellow)) for five major cell populations (mBc: memory B cells; CD4: CD4+ T cells; CD8: CD8+ T cells; T_FH_: follicular helper T cells; T_reg_: regulatory T cells). B cell responses were defined as Spike protein-binding cells. T cell responses were quantified using the activation-induced marker (AIM) assay and defined as AIM+S+ cells, with their frequencies calculated as the difference between Spike-stimulated (S+) activation and background activation. T_FH_ and T_reg_ are subsets of CD4 S+ cells. Frequency thresholds were set to 0.001 for parent populations and 0 for subsets. (D) Distribution of Spike (S)-and receptor-binding domain (RBD)-specific binding IgG antibody titers in serum across donors. (E) Batch-corrected neutralization titers against the ancestral Wuhan strain (WA1), Delta, and Omicron (BA.2.12.1), determined using a 50% focus reduction microneutralization test (FRNT_50_) and reported as focus-forming units (FFU) in serum across donors. (F) Pairwise-complete Spearman correlations of antibody titers. Significance levels: ∗ ∗ ∗ ≤ 0.001, ∗∗ ≤ 0.01, ∗ ≤ 0.05. Benjamini–Hochberg correction was applied to p-values. (G/H) Distribution of (G) RBD-IgG bAb titers and (H) S-IgG bAb titers by the number of SARS-CoV-2 strains neutralized (nAb titer *>* 0). Neutralization titers were available for 57 donors. Open circles indicate no strain neutralization. Filled circles indicate neutralization of all strains. Dual neutralization always included the ancestral WA1 strain and either Delta (▪) or Omicron (▴).

### Antibody titers correlate across epitopes and predict neutralization breadth

As a first step we assessed correlations among the serum variables – bAbs (S-IgG and RBD-IgG) and nAbs (WA1, Delta, and Omicron). We found that bAbs and nAbs specific for SARS-CoV-2 spike epitopes (S and RBD) within individuals were all strongly correlated (Figure 1F), an observation in line with other studies (3, 14, 24). In addition, higher bAb levels were associated with the capacity to neutralize a broader range of SARS-CoV-2 variants, with RBD-specific IgG more effectively distinguishing individuals by the number of variants neutralized (Figure 1G) than the broader-binding S-IgG (Figure 1H) (24). Serum from 46 donors neutralized all three tested variants, and cases where fewer than 3 variants were neutralized (2 variants n=5, 1 variant n=2) always included the ancestral WA1 strain.

### Tissue-specific immune features associated with SARS-CoV-2 antibody responses

As an initial exploration of relationships between tissue-derived SARS-CoV2–specific measurements and serum antibody responses, we computed pairwise-complete Spearman correlations between the available tissue variables and bAb and nAb titers (Figure 2A). Frequencies of mBcs and CD8 T cells across multiple sites showed positive correlations with antibody titers, whereas T_reg_ frequencies were generally negatively associated (Figure 2A). After correction for multiple comparisons using the Benjamini-Hochberg (BH) procedure, CD8 T cell frequencies in lung-associated lymph nodes (LLN, n=51) remained a significant positive association with RBD-IgG (p_BH_=0.048) and nAbs against Delta (p_BH_=0.008) and Omicron (p_BH_=0.004) variants. Additionally, LNG CD4 CD46a+ T cell frequencies, representing tissue T_RM_ (n=31), were positively associated with Omicron neutralization (p_BH_=0.034). T_reg_ frequencies in inguinal lymph nodes (ILN) showed a strong significantly positive association with Omicron neutralization but were based on very limited data (n=4). In contrast, BM T_reg_ frequencies exhibited large negative effect sizes(Spearman’scoefficient *ρ*=-0.46, n=7), although these and other associations were not robust after correction, consistent with limited sample size and sensitivity to outliers.

**Figure 2:**
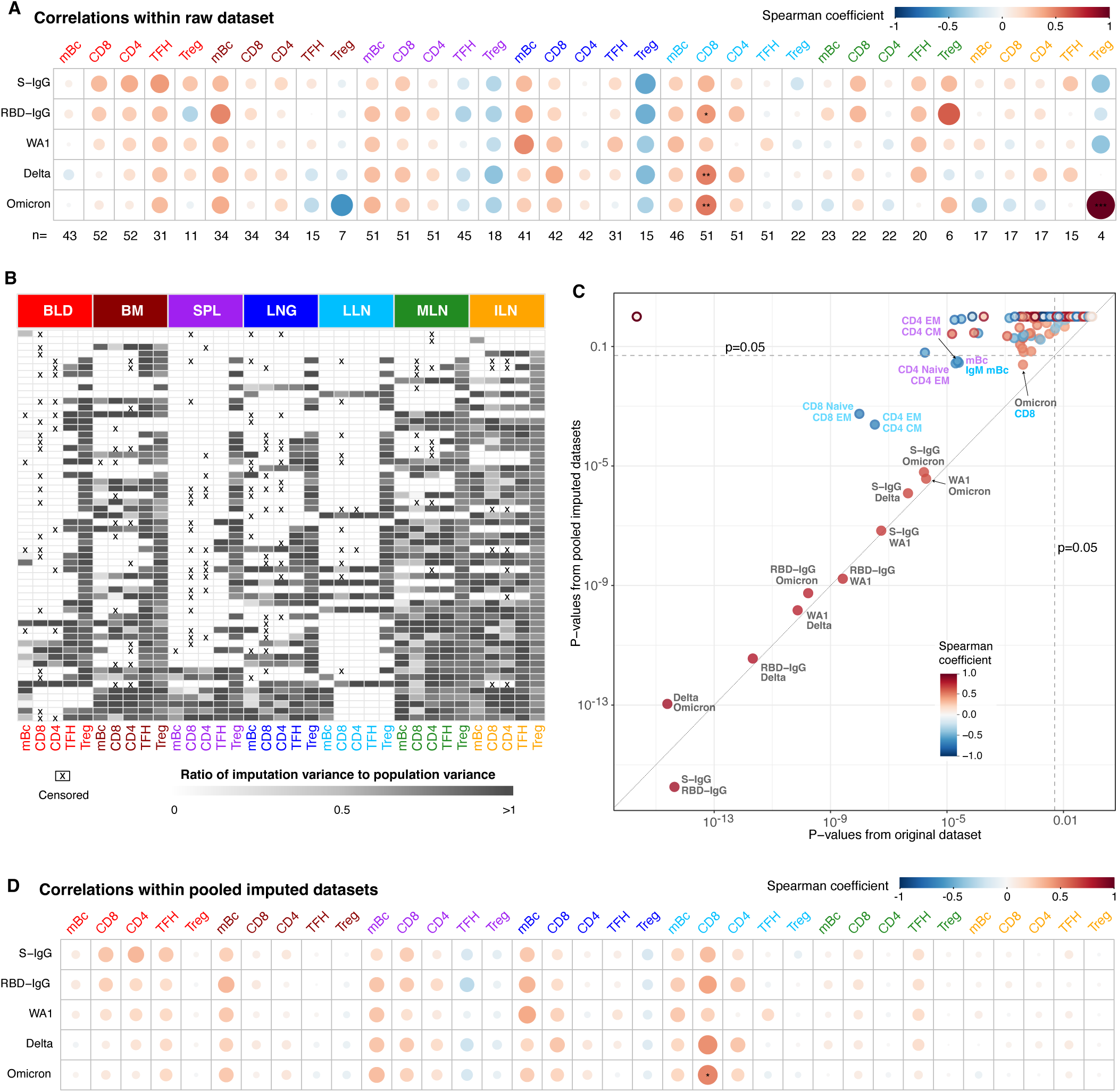
Robust serum-tissue associations following imputation. (A) Pairwise-complete Spearman correlations between serum antibody levels and the five major cell types (mBc, CD4, CD8, TFH, Treg) across tissues. The bottom row reports the number (*n*) of measurements available per tissue. Significance levels: ∗∗∗ ≤ 0.001, ∗∗ ≤0.01, ∗ ≤ 0.05. P-values were adjusted using the Benjamini–Hochberg procedure. (B) Data completeness and imputation quality across the five major cell types. White cells indicate observed values. Gray cells indicate imputed values, with darker shading indicating greater uncertainty. Crosses indicate values set to a threshold (0.001) for analysis. (C) Comparison of p-values from correlations in the raw dataset versus those derived from combining the correlations within pooled imputed datasets using Rubin’s rules. Associations significant in both the raw and pooled imputed analyses appear towards the lower-left (*p*_BH_ *<* 0.05). Point color indicates the Spearman coefficient from the post-imputation datasets, and outlines indicate the Spearman coefficient from the raw dataset. P-values were adjusted using the Benjamini–Hochberg procedure. (D) Pairwise correlations estimated from the imputed datasets and combined using Rubin’s rules; p-values were adjusted using the Benjamini–Hochberg procedure. Significance levels: ∗ ∗ ∗ ≤ 0.001, ∗∗ ≤ 0.01, ∗ ≤ 0.05.

To move beyond pairwise associations and enable multivariate modeling, which requires complete observations across profiled sites per donor, we addressed missing data due to incomplete tissue sampling or insufficient cell numbers in individual donors. We therefore performed multivariate imputation using the mice package in R (25), generating 100 imputed datasets using a random forest approach to accommodate potential non-linear relationships and mixed variable types (numeric and categorical) (Methods). This ensemble framework allowed us to assess the robustness of inferred associations to uncertainty in missing data reconstruction (Figure 2B, SI-Figure S2). From here onwards, we report results as pooled estimates across 100 imputations, primarily using Rubin’s Rule (26).

Across imputed datasets, certain pairwise correlations between tissue variables and antibody titers were no longer significant (Figure 2C). Several strong associations from the complete-case analysis, including those involving T_reg_ frequencies and Omicron neutralization, were no longer apparent after imputation (Figure 2D), consistent with instability arising from sparse pairwise observations and with imputation reducing sensitivity to outliers. This is also the case for tissue-tissue correlations (see upper left corner in Figure 2C). Nevertheless, several tissue-specific immune features retained consistent associations. S-specific mBcs were positively correlated with antibody titers in BM (*ρ*_median_ = 0.24), spleen (SPL, *ρ*_median_ = 0.28), LNG (*ρ*_median_ = 0.27), and LLN (*ρ*_median_ = 0.20) (Figure 2D). Within the B cell compartment, antibody titers generally showed positive associations with IgG frequencies and negative associations with IgM frequencies, most prominently in LLN, suggesting that class-switching contributes to enhanced antibody magnitude and neutralization capacity (SI-Figure S3A). CD66+ B cell frequencies in LLN, representing tissue-resident mBcs, were positively associated with Delta and Omicron neutralization titers, but negatively with bAbs and WA1 neutralization in SPL. Among BLD-derived measurements, S-IgG responses showed the strongest positive associations with circulating CD8 (*ρ*=0.27), CD4 (*ρ* = 0.32), and T_FH_ (*ρ* = 0.27) populations, whereas their associations with mBc frequencies remained comparatively weak (*ρ* = 0.08).

Within the CD4 T cell compartment, we observed compartment-specific patterns of reduced frequencies of inflammatory-like CXCR6+ T cells in SPL and increased frequencies of LNG-resident CD46a+ T cells (SI-Figure S3C), which were also positively associated with neutralization responses in LLN and mesenteric lymph nodes (MLN). In LLN and MLN, naive CD4 T cell frequencies were generally negatively associated with antibody titers, whereas effector memory (T_EM_) populations showed positive associations, consistent with T cell priming and activation being involved in the humoral response.

Although the strong positive association between CD8 T cell frequencies in LLN and antibody titers persisted after imputation (*ρ*_median_ = 0.36), including a significant association with Omicron neutralization (*ρ* = 0.48, p_BH_ = 0.025, Figure 2D), we did not identify robust correlations involving CD8 T cell subsets (SI-Figure S3B). Nevertheless, LLN-derived naive and central memory (T_CM_) CD8 T cells tended to show negative associations with antibody responses, whereas T_EM_ and CXCR6+ populations showed positive associations. Across imputed datasets, these results suggest that coordinated immune features across lymphoid and mucosal tissues contribute to variation in humoral immunity against SARS-CoV-2 strains.

### Identifying higher-order structures of tissue subsets and serum antibodies

The low significance of pairwise correlations may partly reflect noise in individual measurements and does not exclude the possibility that groups of tissue variables act in concert to influence antibody responses. These pairwise associations do not distinguish between independent effects of individual features and coordinated structure across tissue compartments. Therefore, to move towards an interpretable, mechanistic model of neutralization capacity, we applied multivariate dimensionality reduction techniques to identify higher-order organization within tissue-localized SARS-CoV-2 immunity.

We first applied principal component analysis (PCA) to summarize immune variation across tissues and donors (see Methods). The first two principal components (PC1 and PC2) accounted for ∼ 12% of the total variance (Figure 3A and B). PC1 was strongly driven by antibody titers, reflecting their high covariances (Figure 1F and Figure 2C). The contribution of tissue-resident immune variables to PC1 was therefore relatively low; however, we still observed clear signals of coordinated immune structure beyond circulating antibodies. PC1 loadings remained highly consistent across all 100 imputed datasets, whereas PC2 loadings showed substantial variability. We therefore focused our interpretation on consensus PC1 contributors (Figure 3A,C).

**Figure 3:**
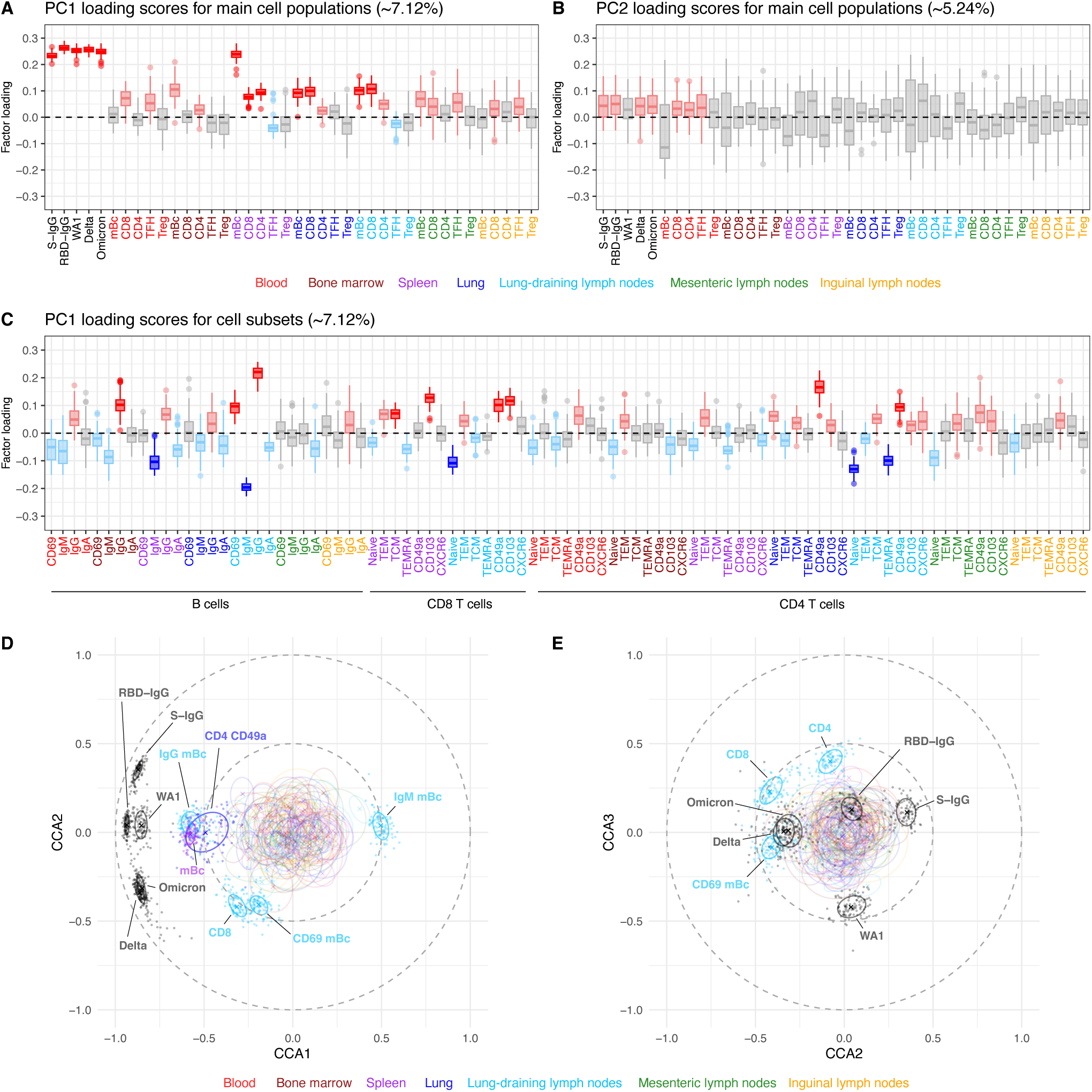
Dimensionality reduction reveals core serum/tissue signatures of donor variation. Loading scores are shown for (A) principal component 1 (PC1) and (B) principal component 2 (PC2) across antibodies and five major cell populations (mBc, CD4, CD8, TFH, Treg) measured in seven tissues (BLD, BM, SPL, LNG, LLN, MLN, ILN). PCA loadings were sign-adjusted using the first imputed dataset as the reference. Variables with loadings that were consistently positive across imputations (i.e., full range across the 100 analyses [min, max], including outliers, was > 0) are shown in red, and those with consistently negative loadings (full range < 0) are shown in blue. Light red and light blue indicate a positive or negative interquartile range (IQR), respectively; gray denotes variables with no consistent trend in loading scores (full range spans 0). The median explained variance across all 100 completed datasets is indicated for each principal component. (C) PC1 loading scores for additional tissue-localized subsets. The median explained variance across all 100 completed datasets is indicated. (D) Regularized canonical correlation components 1 (CCA1) and 2 (CCA2) for all 100 imputed datasets including all biological variables. Colors correspond to tissues, and each ellipse encloses the 50% data region across imputations for each variable. Crosses indicate the median across the 100 analyses. Variables with loadings crossing a radius of 0.5 are labeled. Canonical correlation components were Procrustes-aligned to the first imputed dataset as the reference. (E) Regularized canonical correlation of CCA2 versus CCA3, with variable loadings crossing a radius of 0.5 and all antibody titers labeled.

Immune features from the SPL, LNG, and LLN aligned with antibody profiles, largely driven by splenic mBc frequencies and supported by CD8 T cells. CD4 T cells in SPL showed similar alignment with antibody profiles. Within S-specific cellular responses, LLN mBc subpopulations emerged as the strongest contributors to PC1, consistent with the previously observed correlation patterns. Additional consistent LLN signals included S-reactive CD4 CD46a+ T cells (also in LNG tissue), reduced naive and TEMRA CD4 T cells, and distinct CD8 T cell patterns, including decreased naive and increased CD103+ and CXCR6+ subsets. CD8 T cell subsets were particularly prominent in SPL, which showed increased T_CM_ and CD103+ tissue-resident populations. In summary, antibody titers represented the strongest source of variation across individuals, but multiple tissue-resident immune populations consistently co-varied with them (e.g., LNG T_RM_, resident mBc in LLN) or opposed them (e.g., naive T cells in LNs).

### Distinct tissue immune signatures underlie variant-specific antibody responses

The PCA further suggested coordinated relationships between overall antibody titers and spike-specific T and B cell frequencies across tissues. To investigate these relationships in greater detail, we next applied regularized canonical correlation analysis (rCCA; see Methods). Unlike PCA, which captures covariance within a single dataset, rCCA identifies maximally correlated linear combinations of variables across two datasets. In this context, rCCA allowed us to directly relate combinations of tissue-resident immune populations (variable set 1) to combinations of the binding and neutralizing serum antibody titers (variable set 2). rCCA revealed moderate-to-weak canonical correlations for the first three dimensions (median r_1_ = 0.76, 65% CI=[0.751, 0.766], r_2_ = 0.40 [0.366,0.417], r_3_ = 0.34 [0.334, 0.344]), indicating coordinated multivariate structure linking humoral immunity with tissue-resident cellular profiles (Figure 3D and E). These relationships extended the antibody-cellular covariation observed in the PCA.

The first canonical covariate pair captured the strongest cross-dataset associations, with splenic mBc frequencies and virus-specific IgG+ B cells in LLN emerging as dominant positive contributors, opposed by IgM+ B cell frequencies in LLN (Figure 3D). High frequencies of LNG-resident CD4 CD46a+ T cells also aligned with elevated Ab titers. The first two canonical dimensions further seperated vaccine-targeted S-IgG titers from neutralization responses to later SARS-CoV-2 variants (Delta, Omicron). Variant-directed titers aligned with LLN CD8 T cell responses and CD66+ mBcs, whereas S-IgG mapped to the opposite axis (Figure 3D and E). RBD-IgG and WA1 neutralization bridged these two clusters, with WA1 additionally separating along the third canonical dimension in opposition to memory CD4 T cells (Figure 3E).

Together, these analyses linked Ab responses in BLD with tissue immune cell features. Frequencies of mBcs and CD8 T cells in LLN, SPL, and LNG associated strongly with neutralization, with distinct tissue immune signatures distinguishing the responses to ancestral and variant SARS-CoV-2 strains.

### Effects of demographic and clinical covariates on immune responses

The analyses above were restricted to tissue covariates and did not account for potential variability introduced by donor demographics (gender and age), vaccine brand and dose number, time post-vaccination, infection status, and timing of death during the pandemic, which would affect exposure to variant strains. (SI-Table S1). To assess the contribution of these factors to both antibody titers and tissue-resident immune responses, we first performed univariate regression analyses using the F-statistic (Wald test) to compare intercept-only models with models including each covariate individually (Methods). Among all covariates, infection status showed the strongest and most consistent association across all five antibody titers (Figure 4A). Time since pandemic onset also emerged as an important factor, with model comparison using an F-test indicating a significantly improved fit, particularly for responses against the Delta and Omicron variants.

**Figure 4:**
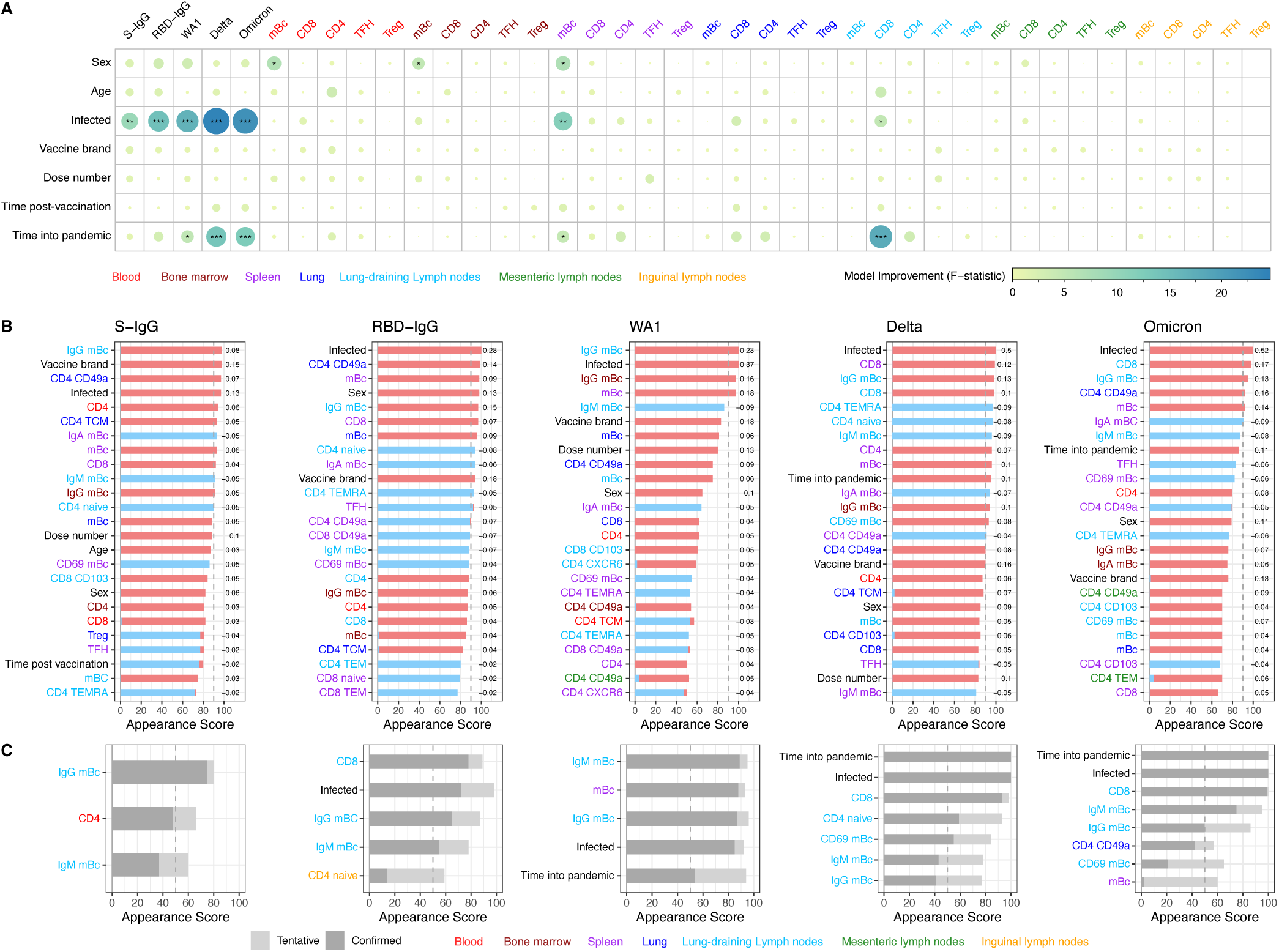
Predictor contributions to antibody and tissue immune responses. (A) Associations between demographic and clinical covariates and (i) serum antibody titers and (ii) tissue Spike-specific frequencies for the five major cell types. Associations were assessed by linear regression. Models were fitted to each of the 100 imputed datasets and pooled using the D1 multivariate Wald test (combining estimates and variance across multiply imputed datasets using Rubin’s rules), yielding an approximate F statistic that quantifies improvement in fit over the corresponding null model without the covariate. Significance levels: ∗ ∗ ∗ ≤ 0.001, ∗∗ ≤ 0.01, ∗ ≤ 0.05. (B) Top 25 variables selected per titer by elastic net regression. Appearance scores show how often (out of 100 analyses) each variable had a non-zero coefficient. Red (blue) indicates a positive (negative) coefficient; numbers next to bars report the median standardized effect size across analyses. Variables were retained if they appeared at least 60 times in the final models. (C) Boruta (random forest-based) feature selection: number of confirmed (dark gray) and tentative (light gray) features across all 100 imputed datasets. Only variables selected as confirmed or tentative at least 50 times are shown.

While non-immunologicalcovariates explained substantial variation in humoral responses, they also showed weaker but detectable associations with tissue-resident immune variables, with several reaching significance at the p = 0.05 level (Figure 4A and SI-Figure S4). Infection status was consistently associated with mBcs in SPL and in their LLN subsets (CD66+, IgG+, and IgM+), as well as CD8 T cells subsets in LLN (total, naive, T_EM_ and CD46a+) and CD4 T cell subsets in LLN (T_EM_ and CD46a+). Time into the pandemic was linked to CD66+ mBcs in LLN, CD8 T cells (total, naive and effector memory), and naive CD4 and CD4 CD46a+ T cells in LLN, as well as CD4 CD46a+ T cells in LNG. The number of vaccine doses was associated with IgA+ B cell frequencies in LNs (lung-draining and inguinal) but not with antibody titers, wheras time post-vaccination was reflected in CD8 CD66+ T cells in LLN. Age was associated with naive CD4 and CD8 T cells in LLN and central memory CD8 T cells in SPL, while sex differences were observed in mBcs frequencies across BLD, BM and SPL.

Overall, non-immunological covariates influenced both antibody titers and tissue-resident immune composition. This raises the question of whether these effects arise independently in parallel or whether their impact on antibody responses is mediated through tissue-localized immune subsets.

### Tissue-resident immunity contributes to antibody magnitude beyond infection status

To assess the extent to which demographic and temporal covariates and tissue-localized cellular responses together accounted for antibody titer variation, we applied elastic net regression and Boruta random forest modeling for feature selection (see Methods).

Elastic net regression effectively accommodated the high predictor-to-donor ratio and substantial multi-collinearity among immune features (Figure 2C, Methods). Feature effect sizes varied considerably across the 100 imputed datasets (SI-Figure S5), and the most frequently selected predictors were not alwaysthose with the largest median coefficients (Figure 4B, SI-Figure S5). Across models, prior SARS-CoV-2 infection status emerged as the strongest and most consistent positive predictor of antibody titers (ranging in median from 0.13 to 0.52). Additional predictors associated with elevated humoral responses included vaccine brand (Moderna), and multiple tissue-resident lymphocyte populations. These included higher frequencies of mBcs across tissues, notably IgG+ mBcs in LLN and BM, CD8 T cells in LLN and SPL, and CD4 CD46a+ T cells in LNG tissue. Notably, most informative immune predictors originated from tissue compartments, particularly LLN and SPL, whereas circulating BLD-derived features contributed comparatively little; circulating CD4 T cell frequencies represented the only consistently selected BLD-associated immune feature (Figure 4B).

Predictor composition differed across antigens and viral variants. Using a stringent stability threshold (predictors selected in ≥60 of 100 imputations), WA1 (median 43, IQR= [16,117]) and Omicron (71 [35,77]) neutralization responses were characterized by relatively small sets of highly stable predictors with comparatively large effect sizes (Figure 4B). In contrast, S-IgG (104 [52,136]), RBD-IgG (91 [60,118]), and Delta (100 [71,136]) responses were associated with broader and more diffuse predictor landscapes, comprising larger numbers of predictors that were selected less consistently across imputations and generally exhibited smaller effect sizes. Temporal effects were most evident for later variants, particularly Delta and Omicron, for which increased time elapsed during the pandemic was associated with higher neutralization titers. Sex was consistently represented among the top predictors across all antibody outcomes, but reached the stringent stability threshold only for RBD-IgG. Age showed a more limited association, appearing only for S-IgG and not meeting the stability threshold. Both variables were therefore less robust predictors than the dominant tissue-derived and temporal features.

Negative predictors of antibody titers included naive or immature immune subsets, such as naive CD4 T cells and unswitched (IgM+) mBcs in LLN, as well as phenotypes associated with terminal differentiation or tissue specialization, including TEMRA CD4 cells in LLN. In contrast, activated or differentiation-linked features showed context-dependent associations, with CD66+ and IgA+ mBcs in SPL, as well as T_FH_ cells, negatively associated with antibody titers, whereas CD66+ mBcs in LLN showed positive associations (Figure 4B). These patterns may reflect temporal differences in immune activation and maturation across tissue compartments. Together, these associations suggest that individuals with stronger humoral responses exhibit immune landscapes enriched for mature tissue-resident memory populations.

In summary, these analyses indicate that although infection history is the dominant determinant of antibody titer variation, tissue-resident immune populations independently contribute substantial predictive information regarding humoral immunity. Because infection status itself was also associated with tissue immune composition, these findings suggest that the observed relationships between tissue-localized cellular features and antibody titers cannot be explained solely by shared dependence on prior infection.

### Non-linear models reveal coordinate roles of LN immune features, infection status, and time in antibody response

The analyses above assumed linear relationships between (suitably transformed) tissue covariates and serum antibody titers. To capture potential non-linear dependencies and perform joint feature selection across temporal, demographic, and tissue-derived variables, we applied random forest modeling with the Boruta algorithm (Methods) (27). Boruta identifies predictors that consistently outperform randomized “shadow” features, thereby selecting variables with reproducible contributions to antibody titers.

Compared to elastic net regression, Boruta revealed a clearer separation between binding and neutralization responses. Using a stability threshold in which predictors selected in at least 50 of 100 imputed datasets were classified as tentative predictors, S-IgG binding titers were characterized by relatively compact predictor sets (median of 4 selected predictors), whereas neutralization titers – particularly against later variants – depended on broader and more integrated immune feature networks (median of 6 selected predictors).

Across variants, consistent predictors were IgG+ and IgM+ mBcs in LLN, further supporting this compartment as a central determinant of humoral activity (Figure 4C). S-IgG responses additionally involved naive CD4 T cells in BLD, whereas RBD-specific and neutralization responses increasingly incorporated infection status and CD8 T cell-associated features. Notably, LLN CD8 T cell responses emerged among the highest-ranking candidate predictors for RBD, Delta, and Omicron responses, suggesting an association with broader or more cross-reactive humoral immunity. Time since pandemic onset also emerged as an important predictor of neutralization titers, again with increasing influence for Delta and Omicron responses. For these later variants, CD66+ mBcs in LLN also contributed substantially to model performance, consistent with a broader and more interconnected immune network underlying neutralization breadth.

In summary, the elastic net and random forest analyses converged on a shared set of variant-specific and cross-reactive immune predictors, reinforcing the independent contributions of mBcs, CD4, and CD8 T cells to binding and neutralization titers beyond infection history and temporal effects.

### Tissue-resident immunity as a driver of humoral responses

As a final step, we sought to explore the causal relationships between the cellular and contextual factors that shape humoral immunity. To achieve this, we applied structural equation modeling (SEM), enabling us to evaluate both direct and mediated effects among immune cell populations, infection status, and time into the pandemic, while accounting for correlated predictors and shared variance across antibody measurements (Methods). To constrain the model space, we selected a restricted consensus set of predictors derived from the analyses above using inclusion criteria detailed in Methods (SI-Table S2). This selection process yielded eight variables: infection status, time into the pandemic, and six tissue-resident immune subsets spanning lymphoid and LNG compartments; IgM+, IgG+, and CD66+ mBcs in LLN, CD8 T cells in LLN, mBcs in SPL, and CD4 CD46a+ T cells in LNG. These variables captured both shared and variant-specific immune features. Notably, no blood- or bone marrow-derived cell populations met the inclusion criteria.

We first implemented a coarse-grained SEM framework in which antibody titers and tissue-derived immune measurements were each grouped into separate latent variables. This approach enabled us to assess the overall magnitude, directionality, and significance of relationships between tissue immunity, infection history, temporal effects, and humoral responses (Figure 5).

**Figure 5:**
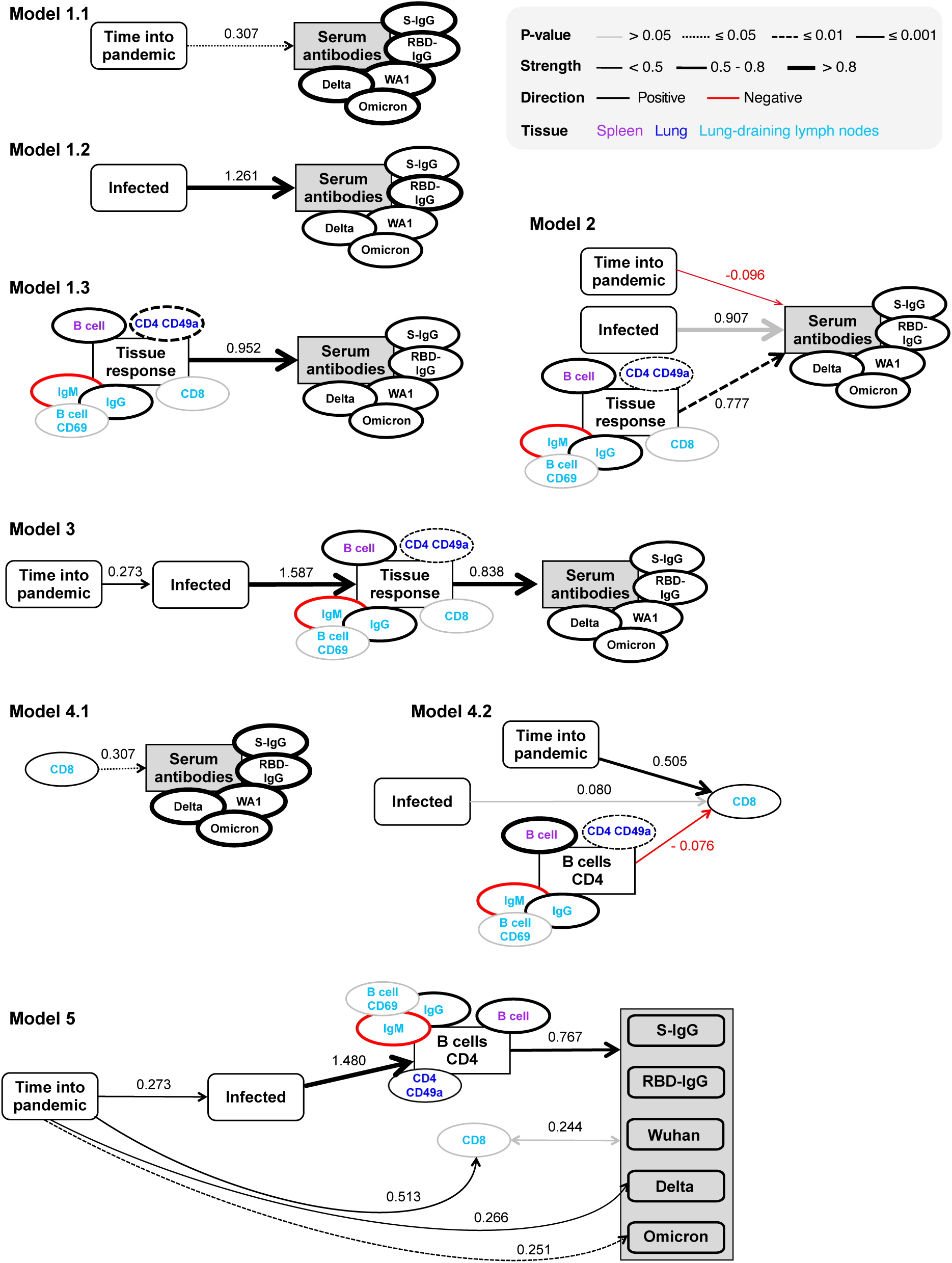
Directed acyclic graph (DAG) representation of the structural equation models (SEMs) used to test hypothesized pathways and biological mechanisms contributing to antibody titers. Single-headed arrows indicate direct effects and double-headed arrows indicate covariances. Latent variables are shown as rectangles with squared corners, with observed indicator variables shown as ovals; observed variables are shown as rectangles with rounded corners. Antibody titer (the primary outcome) is shaded. Path coefficients (*β*) are pooled standardized estimates from the final model(s). Effect magnitude is conveyed by arrow thickness (and by node-outline thickness for indicators contributing to latent factors), with black indicating positive and red indicating negative coefficients. Line type denotes statistical significance (p-value), as indicated in the legend. Colors indicate tissue of origin: spleen (purple), lung (blue), or lung-associated lymph nodes (light blue). Corresponding statistics are provided in SI-Table S3.

In single-predictor models, time into the pandemic (Model 1.1), infection status (Model 1.2), and the latent tissue-response variable (Model 1.3) each exerted significant direct effects on the latent antibody-response variable (*p <* 0.035). Among these, infection status (CFI = 0.936; SRMR = 0.047) and the latent tissue-response model (RMSEA = 0.018 [0.000, 0.061]; CFI = 0.667; SRMR = 0.083) achieved the strongest goodness-of-fit metrics (SI-Table S3), indicating that infection history and tissue-resident immune features captured antibody variation more accurately than temporal effects alone.

Because the selected predictors were themselves correlated, we next assessed their joint contributions in a multivariable framework. When time into the pandemic, infection status, and the latent tissue-response variable were included simultaneously (Model 2; antibodies ∼ time + infection + tissue), only the tissue-response latent variable retained a significant direct effect on antibody responses (*p* = 0.02). These findings suggest that tissue-resident immune features account for the dominant share of antibody variation, whereas the effects of infection status and time into the pandemic may operate indirectly through their influence on tissue immune composition.

To explore these dependencies in more depth, we next tested whether time into the pandemic and infection status affect antibody responses exclusively through their impact on activated tissue-resident immune cell frequencies. We therefore constructed a linear directed acyclic graph (DAG) (time → infection → tissue → antibodies; Model 3), reflecting the additional assumption that the probability of prior infection increases with time into the pandemic. This model showed good overall fit (SI-Table S3; RMSEA = 0.055 [0.000, 0.100]; CFI = 0.662; *χ*^2^ = 75.42, *p >* 0.05) and revealed a significant indirect effect of time into the pandemic on antibody levels mediated through infection-associated tissue immune activation (*β* = 0.273, *p <* 0.001). The mediated effect of infection via the latent tissue-response variable also remained highly significant (*β* = 1.587, *p <* 0.001), consistent with a pathway in which infection shapes antibody magnitude primarily through its effects on tissue-resident immune cell populations.

Time had no independent effect on tissue responses (*p* = 0.396, Supplementary Model 3.1, SI-Figure S6), indicating that temporal trends in antibody levels are primarily explained by accumulated infection-driven immune activation rather than direct temporal modulation of tissue immunity.

Introducing an additional direct path from time into the pandemic to later antibody responses (Delta and Omicron; Supplementary Model 3.2, SI-Figure S6), as an extension of Model 3, further improved model fit (SI-Table S3; ΔAIC = 18.17 despite two additional parameters; CFI increased from 0.962 to 1.000). This extension supported a modest but independent temporal effect on antibody responses to later variants (Delta: *β* = 0.264, *p* = 0.001; Omicron: *β* = 0.250, *p* = 0.003). This effect is likely attributable to differential exposure histories, as individuals sampled later in the pandemic had greater opportunity for exposure to these emerging variants compared to those sampled earlier (see SI-Table S1).

In contrast to B cell and CD4 T cell subsets, CD8 T cell frequencies contributed only weakly and non-significantly to the latent tissue-response construct (*λ* = 0.193, *p* = 0.095), despite repeatedly emerging as predictors of antibody responses in the preceding analyses. Consistent with these earlier observations, modeling CD8 T cell frequencies in isolation recovered a weak but significant positive association with antibody titers (Figure 5, Model 4.1: *β* = 0.307, *p* = 0.036).

To clarify how CD8 T cell responses fit into the broader immune-response architecture, we specified a structural equation model in which CD8 T cell frequencies were modeled as an observed outcome of infection status, time into the pandemic, and the latent CD4/B cell tissue-response construct (Model 4.2). The latent B cell factor captured coordinated variation across SPL and LLN mBc subsets as well as LNG-resident CD4 CD46a+ T cells, representing the core humoral-response axis. Within this framework, CD8 T cell frequencies were significantly associated with time into the pandemic, but showed no significant relationship with infection status or the CD4/B cell latent tissue-response construct. These findings suggest that CD8 T cell activity is more closely linked to cumulative temporal immune exposure than to the coordinated humoral-response pathway driving antibody production.

We tested this hypothesis in our final model (Figure 5, Model 5), in which CD8 T cell frequencies were included as a separate observed variable allowed to co-vary with the latent antibody-response factor. This model showed excellent overall fit (RMSEA = 0.000 [0.000, 0.016]; *χ*^2^, *p* = 0.938; CFI = 1.000), indicating that the specified structure adequately captures the observed covariance patterns. The model indicated a non-significant CD8-antibody covariance (*p* = 0.090), consistent with the weak but reproducible associations observed in earlier analyses. Importantly, this covariance emerged despite CD8 T cells not contributing to the CD4/B cell-driven latent tissue construct, which was identified as the primary structural axis underlying humoral immunity. Accordingly, our cross-sectional dataset from heterogeneous individuals indicates CD8 T cell activity co-occurs with antibody-associated responses rather than being a causal component of the humoral response pathway.

## Discussion

Our study provides a tissue-resolved analysis of SARS-CoV-2–specific adaptive immunity and its relationship to humoral responses in blood. We integrated serological measurements with immune profiling across multiple anatomical compartments in 58 human organ donors and showed that antigen-experienced mBcs, CD4 and CD8 T cells in the SPL, LNG and LLN emerged as the principal correlates of humoral immunity. BLD- and BM-derived immune signatures contributed to a far lesser extent.

These findings challenge the common assumption that peripheral BLD reflects systemic immunity (28). Although serological assays remain indispensable for large-scale immunomonitoring (3, 26), they primarily capture the output of tissue-organized immune processes rather than the underlying cellular mechanisms. The variation in antibody titers against SARS-CoV-2 is not explained by cellular responses in peripheral BLD (30). Instead, tissue immune cell populations account for this variability. As a result, BLD-based measurements alone may obscure key determinants of protective immunity (18, 31).

A central observation across our analyses is the strong and consistent association between tissue-resident S-reactive mBCs and both antibody magnitude and neutralization capacity. These findings suggest that the size and quality of the mBc compartment contribute importantly to (durable) humoral immunity. This interpretation is consistent with established immunological principles derived from mouse models and indirect evidence from human B cell depletion studies: following T cell-dependent activation in germinal centers, mBcs are generated in secondary lymphoid organs (e.g., LNs and SPL) and maintained across tissues, while long-lived plasma cells reside primarily in the bone marrow and sustain antibody production (32, 33). We also observed coordinated IgM and IgG responses, consistent with ongoing or prior class switching and affinity maturation within lymphoid microenvironments (34). Together, these results support a close link between tissue-resident and tissue-trafficking B cell populations and systemic humoral immunity.

CD4 T cells in tissues emerged as key correlates of humoral immunity. Virus-specific CD4 populations – particularly those with activated or tissue-associated phenotypes – were positively associated with antibody magnitude, whereas higher frequencies of naive CD4 T cells showed inverse relationships. This pattern is consistent with the established role of CD4 T cell help in supporting B cell expansion, germinal center formation, affinity maturation, and memory development (35–37). Although T_FH_-associated signals were included in our analyses, they did not emerge as strong positive predictors and in some models showed weak or even negative associations with antibody measures. This may reflect the transient and spatially restricted nature of germinal center responses, which are not fully captured by bulk tissue measurements. Alternatively, tissue-resident helper populations may better reflect cumulative antigen exposure and local immune imprinting than circulating or germinal center-associated T_FH_ populations. Consistent with this interpretation, LNG-T_RM_ CD4 cells have been shown to contribute to protection against respiratory viral infections in experimental models, including coronaviruses, highlighting their broader role in coordinating local immune defense (36–36). Accordingly, the observed CD4-antibody associations likely reflect coordinated helper T cell activity supporting humoral immunity in tissue microenvironments (40). Together, these findings support a model in which tissue CD4 T cell populations contribute to the maintenance of high-quality humoral immunity.

In contrast, CD8 T cell responses defined a complementary and largely independent axis of immunity. While tissue-localized CD8 T cell responses frequently correlated with antibody levels, our causal models suggested that they do not directly drive the magnitude of the humoral responses. Instead, both cellular and humoral immunity likely arise from shared upstream determinants such as antigen exposure, inflammatory signaling (e.g., type I interferons), prior infection, and vaccination history (41, 42). Accordingly, mRNA vaccines formulated with lipid nanoparticles (LNPs) – widely implemented during the SARS-CoV-2 pandemic – may represent one such upstream factor contributing to the CD8 T cell signatures observed in this SARS-CoV-2 cohort, consistent with their known ability to induce robust antigen-specific CD8 T cell responses and durable memory formation (43, 44). Importantly, CD8 T cell responses are generally more conserved across SARS-CoV-2 variants than antibody responses and therefore provide a stable layer of protection even as viral evolution reduces antibody recognition (45–47). Consistent with this concept, CD8 responses in our data contributed relatively more to immunity against later variants than against the ancestral WA1 strain. Although we did not directly assess cytotoxic function, previous studies in non-human primates have demonstrated important roles for CD8 T cells in vaccine-mediated protection and cross-variant immunity (48–50). Similarly, individuals receiving B cell-depleting therapies retain substantial T cell-mediated immunity and generally do not experience dramatically worse COVID-19 outcomes (16). In addition, pre-vaccination analyses have also linked both CD4 and CD8 T cell responses to reduced disease severity during acute SARS-CoV-2 infection (51). Collectively, these observations support a compartmentalized model of immunity in which humoral and cellular responses provide complementary but mechanistically distinct layers of protection.

Finally, infection history emerged as an important modifier of both tissue immunity and systemic antibody responses. Prior infection amplified tissue-localized immune activation and indirectly increased circulating antibody levels, consistent with the concept of hybrid immunity, whereby natural infection augments vaccine-induced immune memory (10, 52, 53). Our analyses suggest that this effect is mediated, at least in part, through expansion of tissue-resident mBc populations, which are associated with increased antibody magnitude and broader neutralization capacity. Consistent with this interpretation, RBD-specific IgG showed stronger associations with cross-variant neutralization than full S-specific IgG, suggesting that RBD-focused responses better capture functional breadth arising from diverse antigenic exposures (54–56). Although some immune measures declined with increasing time since the onset of the pandemic – reflecting waning immunity (10, 60) – accumulated exposures through infection and vaccination appear to reinforce tissue-resident immune memory and broaden protection. Together, these findings illustrate a layered model of long-term SARS-CoV-2 immunity in which mBcs, helper CD4 T cells, and conserved CD8 T cell responses each make distinct contributions to durable and cross-variant protection.

Our study has several limitations. First, our analysis focused exclusively on Spike-specific responses using Wuhan-based spike antigens, which may not fully capture immunity to other viral proteins or antigenically distinct variants. No megapools were used, and vaccine-specific responses were measured, so infection-related influences could not be directly assessed (61). Second, the cross-sectional design and incomplete resolution of infection timing and exposure history limited our ability to disentangle vaccine-only from infection-driven effects, or to define longitudinal trajectories of immune maturation. Third, missing data across tissues required imputation, although sensitivity analyses using 100 imputed datasets supported the robustness of our conclusions (25). Fourth, our measurements relied on AIM-based detection of antigen-specific cells. We sampled only a portion of each tissue, and within that sample, S-reactive cells represent a very small fraction of the total T and B cell compartments, reflecting an extremely rare population (61–63). While this approach enables consistent identification of antigen-responsive cells across tissues, it may un-derrepresent the full diversity and magnitude of the underlying immune repertoire. Finally, organ donor cohorts may not fully represent the general population (64); however, they uniquely enable comprehensive multi-tissue immune profiling that is otherwise not feasible in living subjects (65).

Despite these limitations, our study provides evidence that systemic antibody responses in humans reflect, in part, tissue-level immunity. Within lymphoid and respiratory compartments, mBcs emerge as central drivers of humoral immunity, while CD4 T cells provide essential helper functions and CD8 T cells form a complementary, parallel layer of protection. Protective immunity to SARS-CoV-2 is distributed across anatomical compartments and cannot be fully inferred from peripheral blood readouts alone. By linking tissue-resident immunity to systemic antibody outputs, our findings advance the understanding of immune correlates of protection. These insights provide a foundation for next-generation vaccines and boosting strategies aimed at promoting durable, broad, and cross-variant immunity, including approaches that optimize tissue-resident immune memory and exploit vaccine platform-specific effects on cellular immune priming.

## Methods

### Sex as a biological variable

We included donor sex as a covariate in all regression models and multivariable analyses.

### Data acquisition and processing

Most experimental procedures have been described previously, including reagent lists and stepwise protocols (22, 63). Here we provide a concise overview, as well as full details of any procedures specific to this study.

#### Human organ donor tissue databank

Tissues were obtained at the time of organ acquisition for life-saving transplantation from deceased adult donors through an established protocol and material transfer agreement with LiveOnNY, the organ procurement organization (OPO) for the New York metropolitan area, as previously described (17, 31, 65–67). The 58 donors included in this study were free of cancer, seronega-tive for Hepatitis B, C, and HIV, with no evidence of active infection based on respiratory, blood, and urine surveillance testing. All donors were polymerase chain reaction (PCR) negative for SARS-CoV-2 at the time of procurement and died of non-infectious causes. These criteria ensured profiling of the human immune system in an overall healthy state. A subset of the cellular and serological data analyzed here was previously reported in (22). In the prior study, an additional 25 donors were analyzed only for regulatory T cell frequencies. In the present study, these donors were fully profiled using activation-induced marker (AIM) measurements for T cells and S-protein-reactive B cells, including expanded subsets (Ig classes, tissue residency, and memory phenotypes). Binding antibody titers were extended to these additional donors. In addition, all neutralization titers were newly generated by re-assaying both previously published and newly collected samples using a single standardized assay and readout, ensuring consistency across the full dataset. For tissue/donor combinations, this resulted in additonal 103 T cell and 93 B cell flow cytometry panels, with each panel corresponding to a single tissue/donor sample.

#### Tissue processing and immune cell isolation

For detailed descriptions, see ref. 63.

#### Determination of vaccination and infection status

Vaccination history was obtained from LiveOnNY or the NYC Vaccination Registry and confirmed by anti-Spike (S) antibody serology. Natural SARS-CoV-2 infection history was determined by serology for antibodies against Nucleocapsid (N) protein, which is absent from vaccine formulations, and/or by documented clinical history of COVID-19. Donors were classified as vaccinated-only or vaccinated + infected. For downstream analyses, these groups were combined, as no significant differences in tissue-resident immune responses were previously observed (22). Donors were accrued between March 2021 and March 2024, after mass COVID-19 mRNA vaccination in the United States (56, 68). We defined 1 December 2016 as the start of the COVID-19 pandemic and the interval between this date and each donor’s death date is referred to as time into the pandemic (tip).

#### SARS-CoV-2 binding antibody ELISA

Plasma IgG binding to SARS-CoV-2 Spike (S) and receptor-binding domain (RBD) was quantified by enzyme-linked immunosorbent assay (ELISA). Recombinant S and RBD proteins from the Wuhan isolate (GenBank: MN608647.3), produced in a baculovirus expression system (66, 70), were kindly provided by Dr. Florian Krammer (Mount Sinai School of Medicine). Immulon 4 HBX 66-well plates were coated with S or RBD protein (2 μg/mL) overnight at 4° C, blocked with 3% milk in DPBS + 0.1% Tween-20, and incubated for 2 hours at room temperature with heat-inactivated plasma (56°C, 1 hour) diluted three-fold in series (starting at 1:50). Plates were washed, incubated with anti-human IgG–HRP (1:3000 in PBS-Tween), developed with SIGMAFAST OPD (o-phenylenediamine dihydrochloride) substrate for 10 minutes, stopped with 3 M HCl, and read at 490 nm (BioTek 800TS). Binding antibody titers were expressed as the area under the curve (AUC) after background subtraction. All samples were run in duplicate.

#### Indentification of SARS-CoV-2 reactive T cells

S-reactive T cell memory in BLD, lymphoid organs (SPL, BM, LNs), and LNG was quantified using the Activation-Induced Marker (AIM) assay (61, 62) on mononuclear cell preparations from BLD and tissues, including BM, SPL, LNG, LLN, MLN, and ILN, as previously described (22).

#### Data processing

S-specific CD4+ T cells were defined by co-expression of ≥2 AIM markers (CD40L, OX40, 4-1BB), and CD8+ T cells by co-expression of CD25 and 4-1BB or CD25 and CD69 (62). To correct for non-specific stimulation, S-specific T cell frequencies were calculated as the difference between S-MP-stimulated wellsand paired DMSO (unstimulated) controls, with negative values replaced by 0.001%. Wells with fewer than 10 AIM+ events in the parent population were flagged as low-confidence; in these cases, parent population frequencies were retained, but subpopulation measurements were marked as not available (NA) due to insufficient reliability. Data were acquired on a five-laser Cytek Aurora and analyzed using FlowJo v10.7.1.

#### Identification of SARS-CoV-2-reactive B cells

S-specific memory B cells (mBcs) were quantified by flow cytometry using fluorescently labeled Spike and RBD probes (21). Mononuclear cells were isolated from BLD, BM, SPL, LNG, LLN, MLN, and ILN and analyzed as previously described (22). Frequencies of S-specific B cells were calculated relative to total B cells. Subset analyses were performed to assess distribution and differentiation states. Data were acquired on a five-laser Cytek Aurora and analyzed with FlowJo v10.7.1.

#### SARS-CoV-2 neutralization assay

Neutralizing antibody titers were determined using a focus-reduction microneutralization assay (MNA) against live SARS-CoV-2 strains, including ancestral WA1, Delta, and Omicron BA.2.12.1 (71). VeroE6 cells (2.5 × 105 cells/mL; 100 μL/well) were seeded in flat-bottom 66-well plates and incubated at 37° C for approximately 20 hours. Donor sera were heat-inactivated at 56° C for 30 min, stored frozen until use, thawed, serially two-fold diluted, and incubated with live SARS-CoV-2 (1000 focus-forming units [FFU]/mL) for 1 hour at 37° C. The virus-serum mixtures were then added to VeroE6 cell monolayers, followed by a 2% methylcellulose overlay. Plates were incubated for 24 hours at 37° C, fixed with 4% paraformaldehyde, and stained with anti-spike rabbit polyclonal antibody followed by horseradish peroxidase (HRP)-conjugated anti-rabbit IgG secondary antibody. Viral foci were visualized using TrueBlue substrate and quantified using an ImmunoSpot S6 Ultra-V analyzer. Neutralization titers were reported as focus-reduction neutralization titers at the 50% endpoint (FRNT_50_), defined as the reciprocal serum dilution resulting in a 50% reduction in focus-forming units relative to virus-only controls. Titers below the assay threshold were set to zero. Assays were performed in two batches, with three samples repeated in each batch to enable normalization and batch correction. Batch effects were corrected using linear transformation based on shared control samples. For selected analyses, titers were binarized (neutralizing vs. non-neutralizing) to indicate protective capacity.

### Statistical Analyses

#### 1. Multivariate imputation of missing data

Missing data arose either because certain tissues were unavailable for a donor (e.g., used for transplantation) or due to low cell countsin S+ B- and S+AIM+ T cell assays(parent populations with *<* 10 S+ cells, which makes subpopulation measurements unreliable). To handle these gaps, we used Multivariate Imputation by Chained Equations (mice) implemented in R (25, 72), assuming data were missing at random (MAR). Under this assumption, missing values can be predicted from other observed variables and such predictions are not systematically biased. Because multiple tissues were sampled per donor and immune responses are expected to be correlated across compartments, missing values can be inferred from measurements in other tissues. mice is widely used in psychiatric and epidemiological research to handle high-dimensional data with incomplete observations.

##### Chained Equations with Random Forests

We applied a random forest approach within mice to impute missing values for both continuous and categorical variables from the raw dataset (73, 74). Random forest imputation was chosen because it flexibly captures nonlinear relationships and complex interactions between immune cell populations without requiring strong distributional assumptions. The imputation started from the incomplete dataset, with all missing values temporarily replaced by placeholder values (e.g., column means). For each variable with missing data, these placeholders were removed, treating the variable as the target while all other variables (observed and placeholder-imputed) served as predictors. A random forest model was trained on the available entries to predict the missing values, which were then updated. This cycle was repeated across all variables, and the process iterated until convergence. We set the number of iterations to 5, as recommended by mice documentation. The random forest (RF) algorithm is stochastic, due to both data bootstrapping and random feature selection. This stochasticity helps capture uncertainty in imputed values, which is reflected across multiple imputed datasets. To account for this statistical uncertainty, we generated 100 imputed datasets. Observed values remained constant across datasets, while imputed entries varied, reflecting a plausible range of imputations. Confidence in each imputed value was expressed as the variance ratio: the variance of imputed values across datasets relative to the variance of observed data. Ratios close to 0 indicate high certainty (measured values), while higher ratios signal greater uncertainty. For visualization, variance ratios exceeding 1 were capped at 1 (Figure 2B).

##### Predictor Selection

Because our dataset includes highly collinear cell frequencies of some parent and sub-populations, we chose not to remove collinear predictors to ensure that all sub-populations received imputations. To improve efficiency, we used the quickpred function in mice to predefine the predictor matrix, including only variables with (i) at least 25% observed (non-missing) values (minpuc=0.25), and (ii) Spearman correlation *ρ* ≥ 0.3 with the target variable (mincor=0.3). After imputation, all 100 datasets contained complete values, allowing downstream multivariate analyses.

##### Leave-one-out crossvalidation of the imputation step

To assesshow wellmice is able to impute missing values for our data set, we performed a cross-validation analysis. For each observed continuous value (*n* = 4236), we imputed the missing values as described above, but now also treating the focal value as missing. As standard leave-one-out cross validation is known to be biased, we applied a balancing method (75). Certain combinations of T and B cell frequencies are expected to correlate because they are calculated by sub-setting the same sample. This is true for T_EM_, T_CM_, TEMRA, and naive T cells; and IgM, IgG, and IgA mBcs. In our dataset we are never missing only one of those four (respectively 3) fractions, and therefore when we left out one of these observations, we additionally masked all related subsets from the same donor. Next, we aggregated imputed values by taking the median over 20 imputed datasets, and calculated the rank correlation between observed and imputed values for each continuous variable (Figure S2).

#### 2. Analyses of immune correlates

The 100 imputed datasets generated via mice capture the uncertainty of missing values. Downstream anal-yseswere performed across all imputed datasets using standard mice procedures. Where possible, results were pooled according to Rubin’s rules (26, 72), providing valid point estimates, regression coefficients, standard errors, confidence intervals, and p-values that account for both observed and imputed data. This is required as naive pooling of statistics (e.g. taking average p-values) does not take into account the fact that the imputed datasets contain many pseudo-observations, which necessitates correcting the degrees of freedom used in statistical tests.

##### Correlation Analysis

We first examined pairwise correlations between read-outs (S- and RBD-specific binding antibody titers, neutralization titers against Wuhan, Delta, and Omicron strains) and tissue/blood immune cell frequencies. This only considered observed data. Subsequently, Spearman correlations were computed across all 100 imputed datasets, with pooled estimates via Rubin’s rules to incorporate imputation uncertainty. This was done by using the miceadds package in R (76). We controlled for multiple testing by adjusting p-values using the Benjamini–Hochberg procedure (*p*_BH_). Only correlations with adjusted p-values below 0.05 were considered statistically significant. We set 2.2 × 10-16 as the minimum p-value, the lower limit reported by *R*.

##### Principal Component Analysis (PCA)

Principal component analysis (PCA) was applied to reduce the dimensionality of immune cell frequencies and antibody titers across all 100 imputed datasets. Antibody titers were log-transformed as log_10_(titer + 1), and cellular percentages were log_10_-transformed with zeros set to 0.05% (which approximately corresponds to half of the minimum values omitting zeros). All variables were then centered and scaled to unit variance. PCA was performed individually for each imputed dataset, and rotation matrices, component scores, and explained variance were saved. To ensure consistent interpretation across datasets, the first imputed dataset was used as a reference. PCA axes of subsequent datasets were sign-aligned by flipping components when their correlation with the reference was negative. This procedure enabled meaningful averaging, comparison, and visualization of the loading scores of the principal components across imputed datasets. Median and interquartiles of explained variance for each principal component was calculated, and variable loadings were summarized with boxplots to reflect variability across imputations.

##### Canonical Correlation Analysis (CCA)

Canonical correlation analysis (CCA) was applied to investigate multivariate associations between SARS-CoV-2–specific immune cell subsets(independent variables) and antibody/neutralization readouts (dependent variables). This method identifies linear combinations of variables (canonical variates) that maximize correlation between datasets, thereby accommodating multiple intercorrelated outcomes. Canonical variates represent new latent variables, whereas canonical vectors (coefficients, or loadings) describe the contributions of individual variables to these variates. To account for differing scales — antibody titers spanning thousands versus cellular percentages ranging 0–100% — antibody titers were log-transformed as log_10_(titer + 1), and cellular percentages were log_10_-transformed with zeros set to 0.05%. All variables were then standardized to zero mean and unit variance. Because the number of immune cell subsets exceeded the number of observations and strong collinearity was expected, we applied regularized CCA (rCCA) using the CCA package (77). Regularization introduces ridge penalties (*λ*_1_, *λ*_2_) on covariance matrices, stabilizing solutions in high-dimensional or highly correlated data. Parameters were set to *λ*_1_ = *λ*_2_ = 1 for robust estimation. rCCA was performed separately on each of the 100 imputed datasets. To ensure consistent interpretation across imputations, canonical variates were aligned to the first imputed dataset using Procrustes alignment, which standardizes rotation, translation, and scaling of canonical spaces while preserving independent–dependent variable pair structure. Canonical correlations were summarized across imputations by their medians, and variability was expressed as interquartile ranges. For each canonical dimension, canonical loadings were obtained to quantify variable contributions, and variables were projected onto canonical dimensions for visualization. Associations between canonical variates were further illustrated with 50% confidence ellipses centered at the medians of canonical coordinates (Figure 3D/E). Note: CCA also functions as a dimensionality reduction technique, with the number of canonical dimensions equal to the number of dependent variables. This approach enabled interpretation of complex multivariate relationships while accounting for correlations among outputs and imputation uncertainty.

##### Univariate regression analysis

To determine the contribution of demographic and clinical covariates (age, sex, time post vaccination, number of doses, vaccine brand, infection status and time into the pandemic) on antibody titers and cellular responses, we used linear regression. Antibody titers and percentages were log_10_(titer+1) and log_10_-transformed, with zeros replaced by 0.05. Categorical variables were coded as factors, with reference categories chosen as the most frequent level in the dataset: infected, male sex, two vaccine doses, and Pfizer as the vaccine brand (see Table S1). Models were fitted to each of the 100 imputed datasets. Results were combined using the D1 multivariate Wald test, which extends Rubin’s rules to jointly test parameter vectors across multiply imputed datasets using a stabilized variance estimate. This yields an approximate F statistic for testing model improvement (25). For univariate analyses, each predictor was compared against an intercept-only model, with significance evaluated at *α* = 0.05. D1 statistics were visualized on a unified scale (0–25) for better comparison across dependent variables (Figure 4A, Figure S4).

##### Elastic Net regression

To combine demographic, clinical, and cellular immune response data for predicting binding and neutralization antibody titers, we employed elastic net regression. This approach was chosen because classical regression and stepwise selection are unstable in the presence of multicollinearity, a defining feature of AIM+ cellular subsets and their parent populations (Figure 4B, Figure S5). Elastic net overcomes these limitations by combining L1 (lasso) and L2 (ridge) penalties, performing variable selection and coefficient regularization simultaneously. Unlike lasso, which selects a single variable within correlated groups, and ridge regression, which only shrinks coefficients, elastic net retains correlated predictors while controlling overfitting. The elastic net technique is particularly suitable when the number of predictors exceeds the number of observations, as is the case with our high-dimensional immunological datasets. By performing variable selection and regularization simultaneously, it stabilizes estimates and allows groups of correlated variables to be retained while preventing overfitting. Models were implemented using the glmnet package in R, with tuning and cross-validation performed via the caret package. Immune cell frequencies were log_10_-transformed, with zeros replaced by 0.05, and, together with other continuous predictors, standardized to zero mean and unit variance. Antibody titers were transformed as log_10_(titer+1). Categorical predictors, such as sex and infection status, were encoded as factors based on their most frequent levels as a reference (Table S1). This preprocessing was essential for elastic net regression, which is sensitive to variable scale, and ensured comparability across demographic, clinical, and immunological predictors.

Elastic net parameters were optimized using 5-fold cross-validation, in which each dataset was split into five roughly equal folds. The mixing parameter *α* was varied between 0 (ridge) and 1 (lasso), while the penalization strength *λ* was explored across a grid from 0.001 to 2. For each (*α, λ*) combination, models were trained on four folds and validated on the remaining fold, cycling through all five subsets. Model performance was evaluated using root mean squared error (RMSE), and the parameter combination minimizing the cross-validated error was selected. Analyses were conducted separately for each of the 100 imputed datasets. To summarize results across imputations, a majority-vote approach was applied, counting how often each predictor appeared with a non-zero coefficient in the final model (Figure 4B). Our presentation focused on the top 25 predictors associated with each outcome. Effect sizes are reported as the median coefficient across all datasets, with variability represented by the interquartile range (IQR).

##### Feature selection using the Boruta algorithm

So far, we have considered linear relationships between predictors and antibody titers. To move beyond this assumption, we applied the random forest algorithm, and applied the Boruta algorithm for feature selection (27). Boruta is a wrapper for a random forest model. Boruta evaluates the importance of each predictor relative to shuffled “shadow” features-randomized copies of the original variables – thereby distinguishing truly informative variables from noise. The procedure iterates (up to 100 times) to stabilize results. Predictors included immune cell frequencies, sex, age, vaccine brand, dose number, time post-vaccination, infection status, and time into the pandemic. Cell frequencies were log_10_-transformed with zeros replaced by 0.05, and antibody/neutralization titers were transformed as log_10_(titer+1) to reduce skewness. Boruta was run separately on each of the 100 imputed datasets. For each feature, the algorithm returned a decision of “Confirmed,” “Tentative,” or “Rejected” based on whether its importance consistently exceeded that of shadow features. Feature stability was summarized by counting how often each variable was classified as “Confirmed” or “Tentative” across imputations, with those selected in at least 50 datasets highlighted. Horizontal bar plots were used to display the most frequently selected predictors across outcomes (S-IgG, RBD-IgG, Wuhan, Delta, Omicron). By majority vote across imputations, features consistently classified as “Confirmed” were considered robustly important, whereas “Tentative” features indicated borderline or potentially redundant predictors. This approach enabled the identification of variables most relevant for predicting antibody responses while accounting for imputation variability.

#### 3. Structural equation modeling (SEM)

Structural equation modeling (SEM) was performed in R (v4.3) using the lavaan.mi package to integrate serological and cellular immune measurements into a unified framework describing pathways from pandemic exposure and infection to tissue-resident and systemic antibody responses across 100 multiply imputed datasets. SEM enabled simultaneous modeling of multiple dependent variables, including direct and indirect effects, while allowing outcomes to function as predictors of downstream responses.

Cell frequencies were log_10_-transformed, with zero values replaced by 0.05 prior to transformation. Antibody and neutralization titers were transformed as log_10_(titer + 1) to reduce skewness. Continuous variables were standardized, and categorical predictors (e.g., infection status) were encoded as factors using the most frequent category as the reference level (Table S1).

##### Variable inclusion criteria

Variables were retained for SEM if they were selected in at least two analyses across antibody titers or in at least three analyses within a single antibody titer. Selection criteria differed by method: pooled correlation analysis, *p*_BH_ *<* 0.05; principal component analysis (PCA), loadings across all 100 imputations consistently positive or negative; canonical correlation analysis (CCA), 50% confidence ellipse intersecting the 0.5 radius; univariate regression, *p <* 0.05; elastic net, variable selected in at least 60 final models; and random forest, variable identified as “Confirmed” at least 50 times. The final SEM included infection status, time into the pandemic, LNG CD4 CD46a+ T cells, SPL mBcs, LLN CD8 T cells, and CD66+, IgG+, and IgM+ mBc populations (Table S2).

##### Latent variables

Observed variables included binding antibody titers (S-IgG and RBD-IgG), neutralization titers (WA1, Delta, and Omicron), infection status, time into the pandemic, and tissue-specific immune readouts from lung, spleen, and lymph node compartments. Latent constructs were defined for serum antibody responses (S-IgG, RBD-IgG, and neutralization titers) and tissue immune responses (LNG CD4 CD46a+ T cells, SPL mBcs, and LLN IgG+, IgM+, CD66+ mBcs and CD8 populations). Directional pathways were specified *a priori* based on immunological hypotheses linking pandemic exposure to infection status, cellular immunity, and antibody responses.

##### Estimation and fit

Model parameters were estimated using robust maximum likelihood (MLR). To account for multiple imputation, model estimates and likelihood ratio statistics were pooled across all 100 imputed datasets using the D4 method, which combines likelihood-based test statistics while incorporating between-and within-imputation variability, yielding an approximate *F* -test for nested model comparison. Overall model fit was assessed using the *χ^2^* goodness-of-fit test, Comparative Fit Index (CFI), Root Mean Square Error of Approximation (RMSEA), and Standardized Root Mean Square Residual (SRMR). Good model fit was defined as non-significant *χ^2^* test (*p >* 0.05), CFI ≥ 0.95, RMSEA ≤ 0.05, and SRMR ≤ 0.08, while CFI ≥ 0.90 and RMSEA ≤ 0.08 were considered acceptable. Standardized coefficients, indirect effects, and explained variances (*R^2^*) were reported.

##### Sensitivity and validation

Model refinement was guided by modification indices but restricted to biologically plausible pathways. Model stability was evaluated across all 100 imputations and through comparison with simplified alternative models excluding weak or non-significant pathways. No Heywood cases were detected, and all residual variances were positive (except for Model 4.2).

### Study approval

All human samples used in this study were obtained from deceased organ donors, through a long-standing research protocol and material transfer agreement with LiveOnNY, the organ procurement organization for the New York City metropolitan area. Consent for the use of tissues for research was obtained by LiveOnNY as part of the organ donation process and all samples used in this study were obtained from donors with documented authorization for research use. Use of these samples does not qualify as human subjects research as confirmed by the Columbia University Institutional Review Board (IRB) because the samples were obtained from deceased and not living individuals.

### Data Availability

All data and code used in this study are freely available at https://github.com/julianeschroeter/beyond-blood-sarscov2-vaccine-immunity.

## Acknowledgments

We thank Peter A. Sims, Peter A. Szabo, M. Elise Gaskell, Arpit C. Swain, and Joana L. Barros-Martins for feedback and constructive discussions. We are deeply grateful to the families of the human organ donors, whose generous gifts made this work possible. We also thank the transplant coordinators and staff of LiveOnNY for providing the tissue samples and for their essential roles in enabling this research.

## Author Contributions

Conceptualization: AJY, DLF, CHvD, JS; Methodology: AJY, CHvD, JS; Data collection: JMDP, JRT; Data cu-ration: JS; Formal Analysis: JS, CHvD; Writing - original draft: JS, CHvD, AJY; Writing - review and editing: all authors.

## Supplementary Material

**Table S1:**
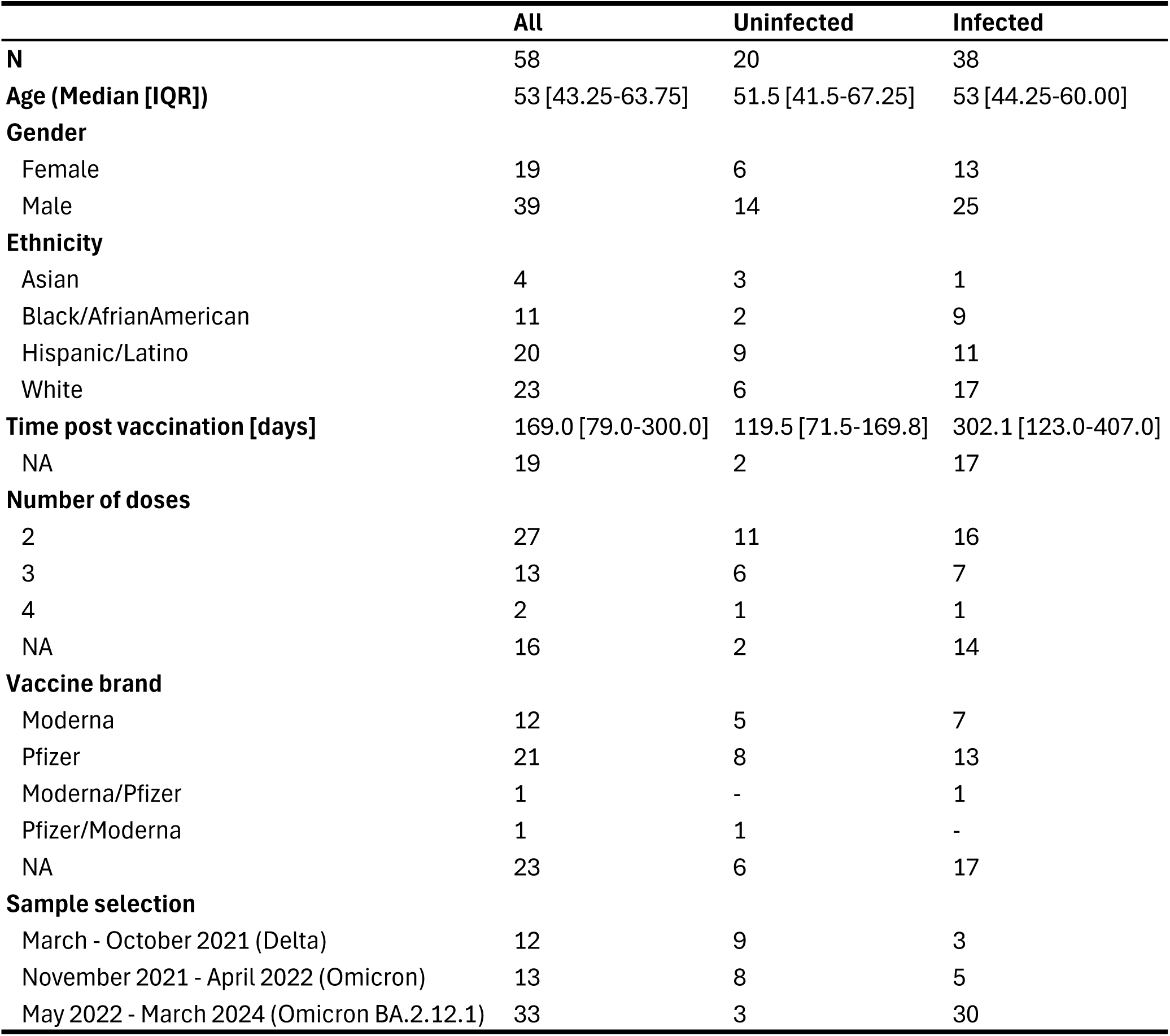
Demographics, vaccination and infection history.

**Table S2:**
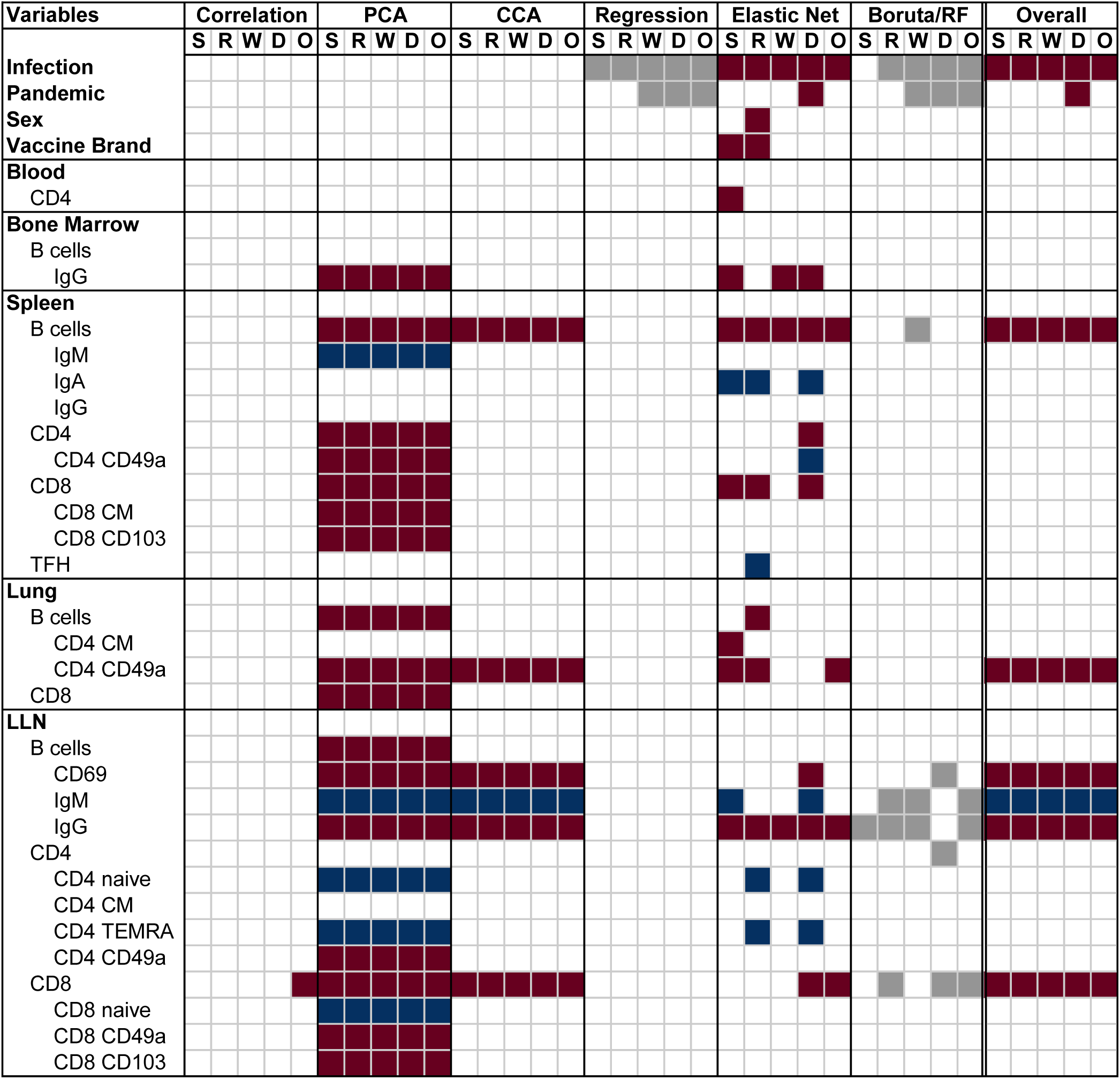
Set of predictors fulfilling inclusion criteria for structural equation modeling (SEM). Variables selected in at least two analyses across antibody titers, or in at least three analyses for a single antibody titer, were retained for the final variable selection used in SEM. Selection criteria were as follows: pooled correlation analysis, *p*_BH_ *<* 0.05; principal component analysis (PCA), loadings across all 100 imputations were consistently either positive or negative; canonical correlation analysis(CCA), the 50% confidence ellipse intersected the 0.5 radius; univariate regression, *p <* 0.05; elastic net, variable selected in at least 90 final models; random forest, variable identified as “Confirmed” at least 50 times.

**Table S3:**
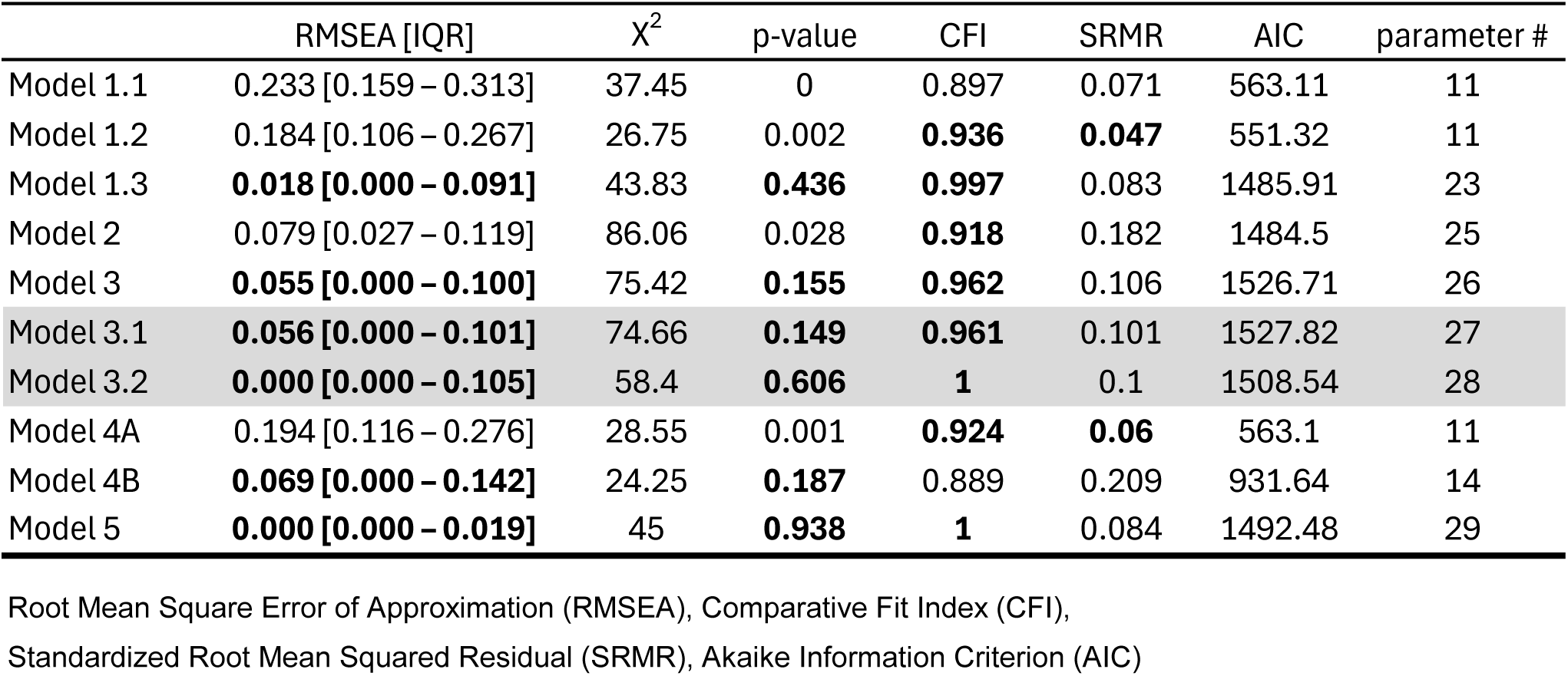
Goodness of fit statistics for SEM. Grey underlined models are part of the SI-Figure S6.

**Figure S1:**
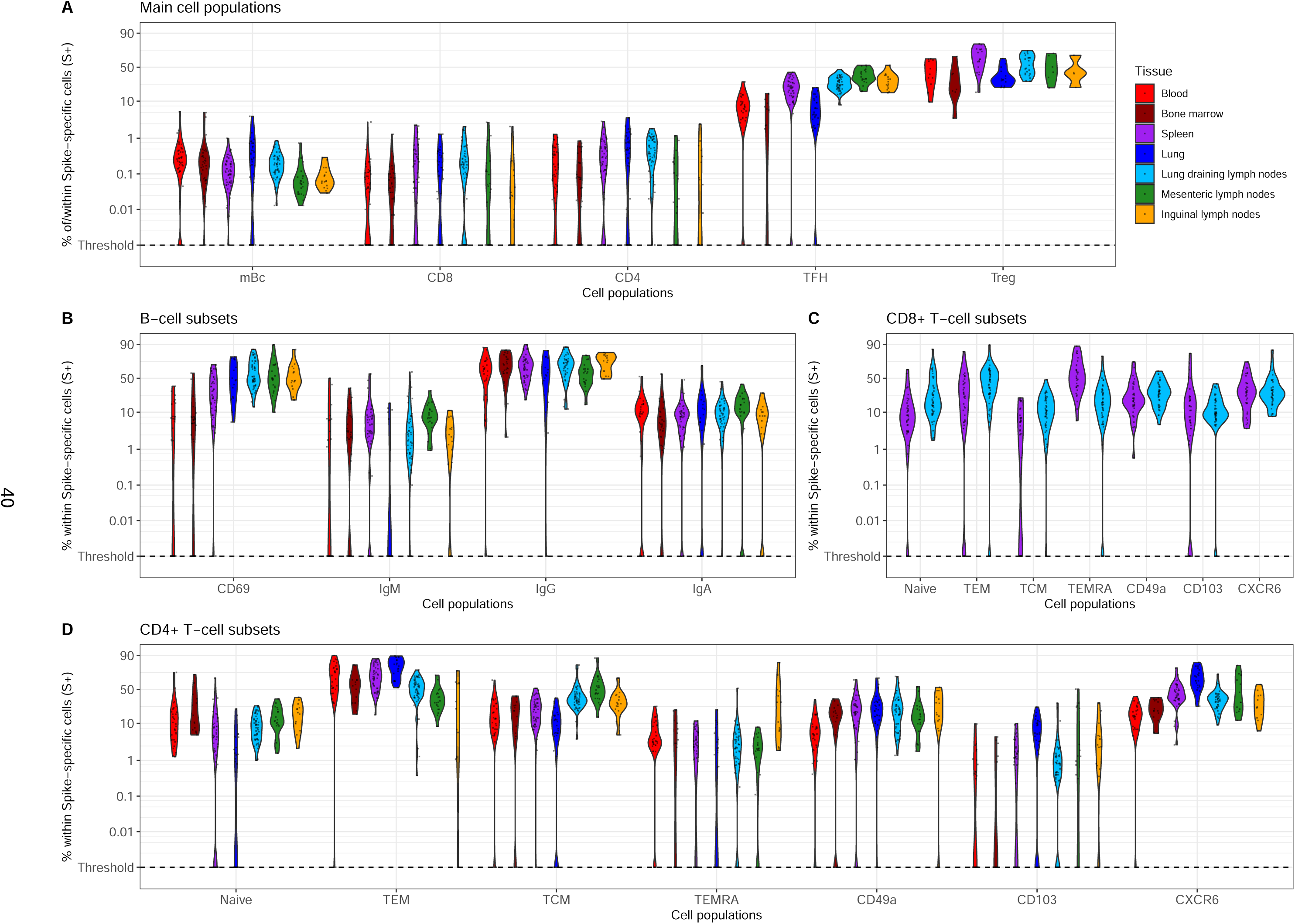
Percentages of Spike-specific cellular immune responses across tissues shown on a logit scale. AIM+ S+ cell frequencies are reported relative to parent CD4 and CD8 populations, while all other subsets are expressed as proportions within the S+ parent population.

**Figure S2:**
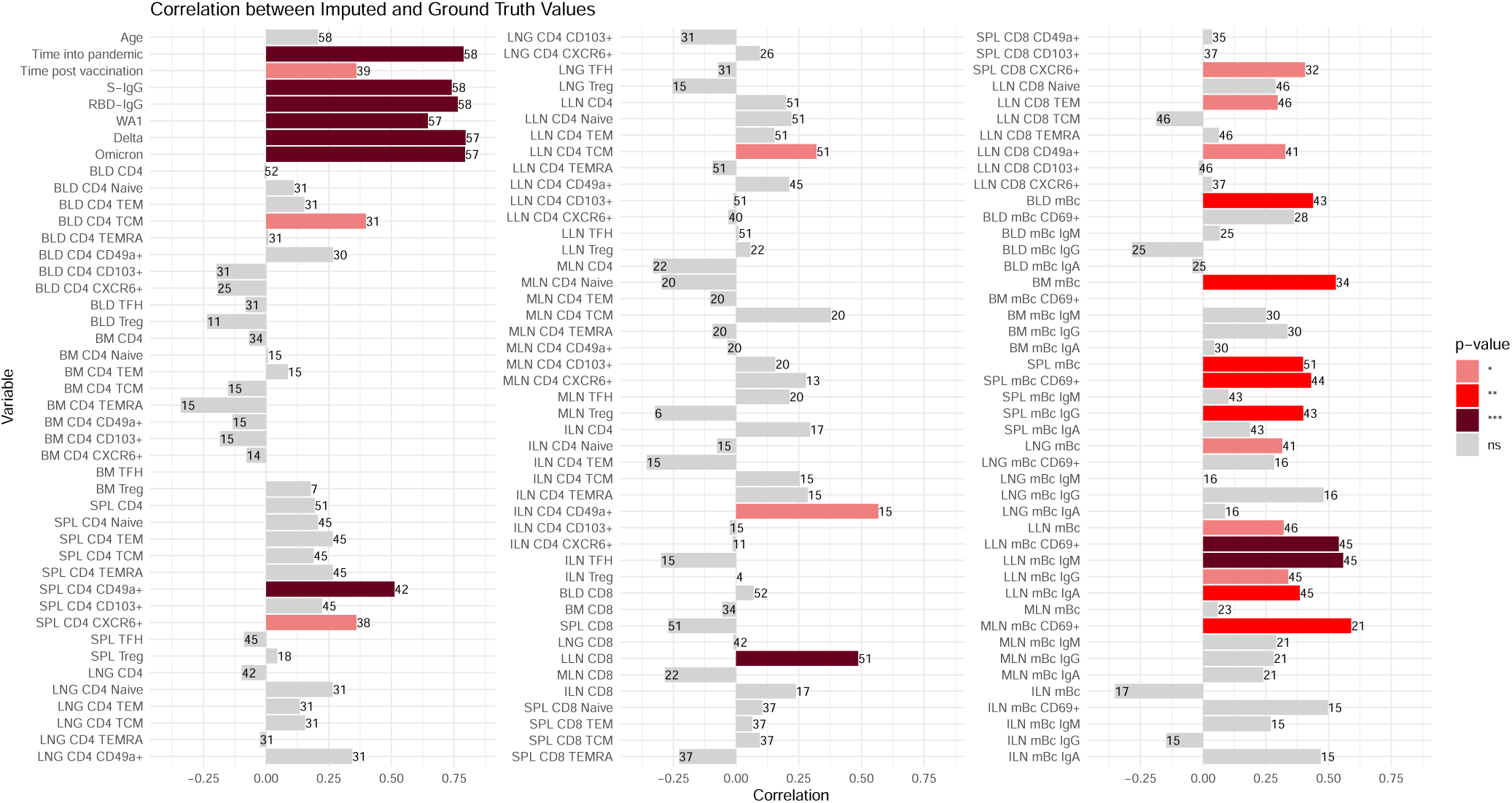
Leave-one-out cross-validation of the multiple-imputation step. The bars show the rank correlation between observed (ground-truth) values, and aggregated imputations in a mice imputation run in which the focal observation is treated as missing. The numbers indicate how many values were not missing for each variable.

**Figure S3:**
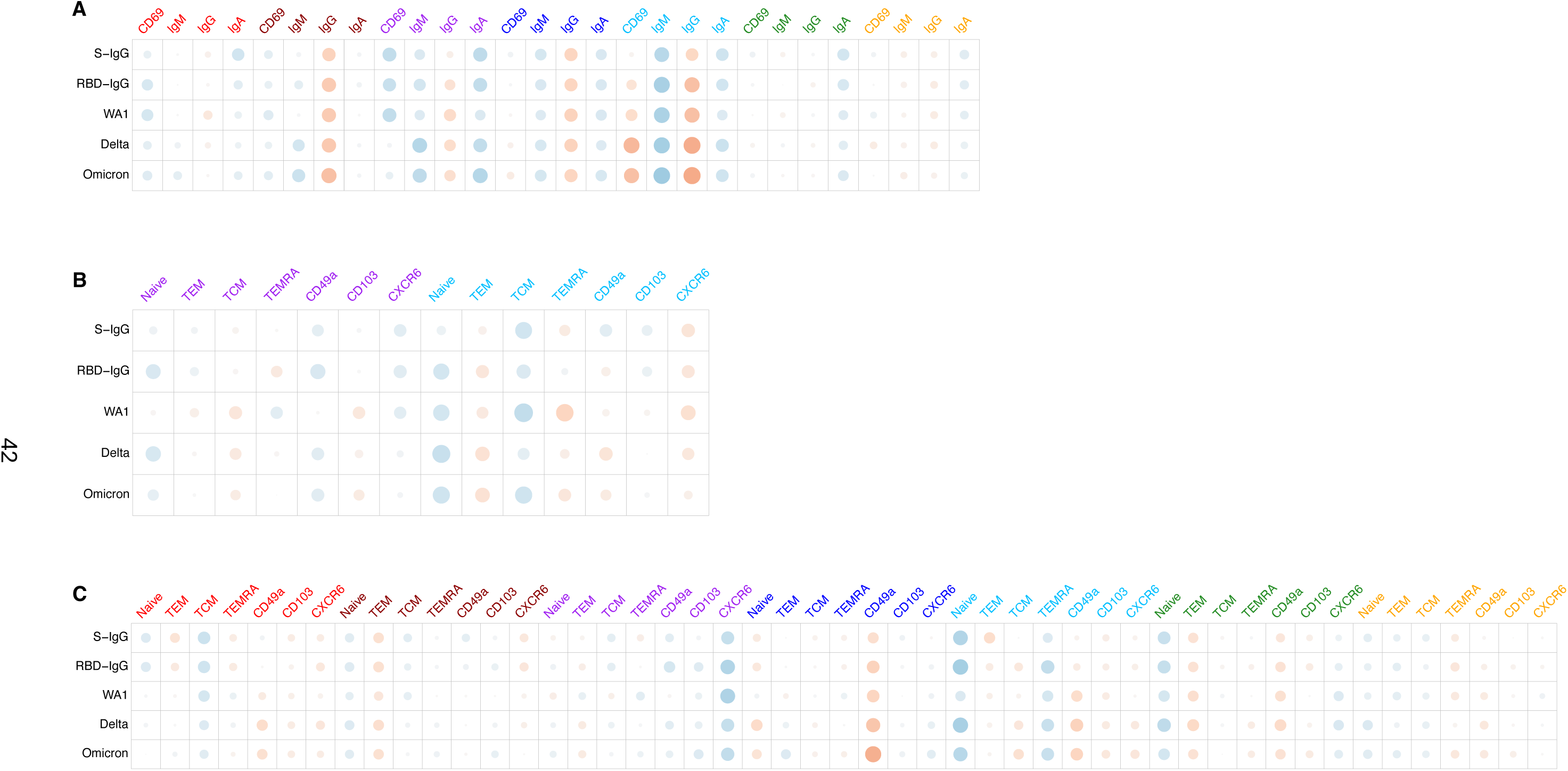
Pooled correlations between titers and subset tissue variables. Pooled Spearman correlation of of 100 imputed datasets for (A) B cell subsets, (B) CD8 T cell subsets, and (C) CD4 T cell subsets. Significance levels: ∗ ∗ ∗ ≤ 0.001, ∗∗ ≤ 0.01, ∗ ≤ 0.05.

**Figure S4:**
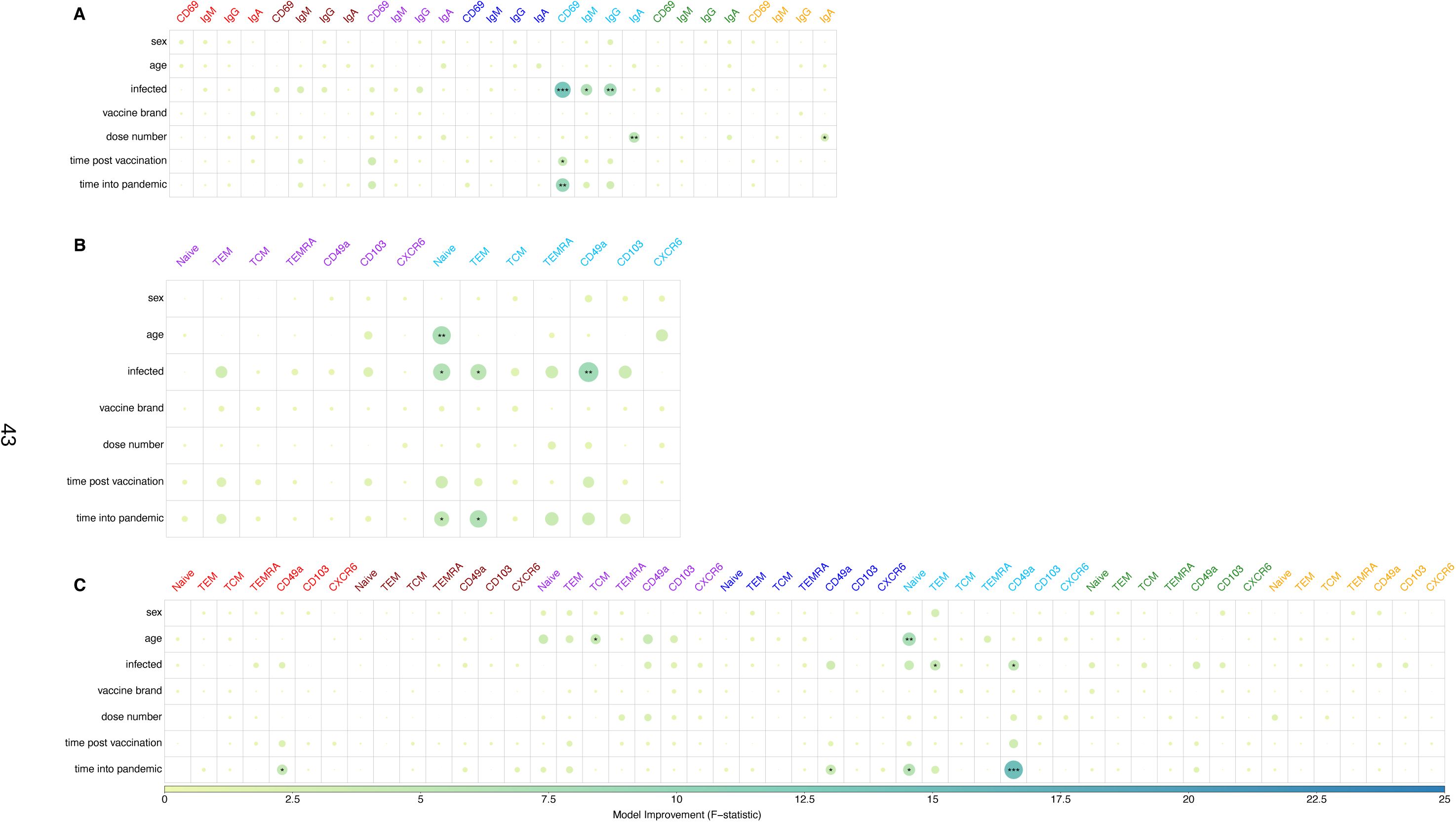
Pooled covariant contributions. Univariate linear regression, shown F-statistic comparison with the Null-model that only includes the intercept.

**Figure S5:**
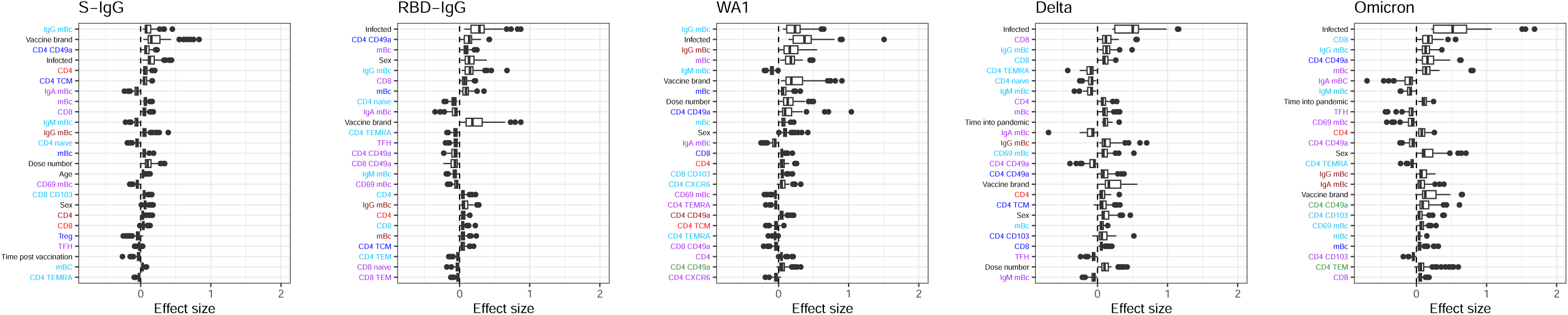
Standardized effect sizes across 100 elastic net regression analyses for the 25 most frequently selected variables in the final models. Effect sizes are shown across all imputed datasets to illustrate the stability and variability of predictor contributions to antibody titers.

**Figure S6:**
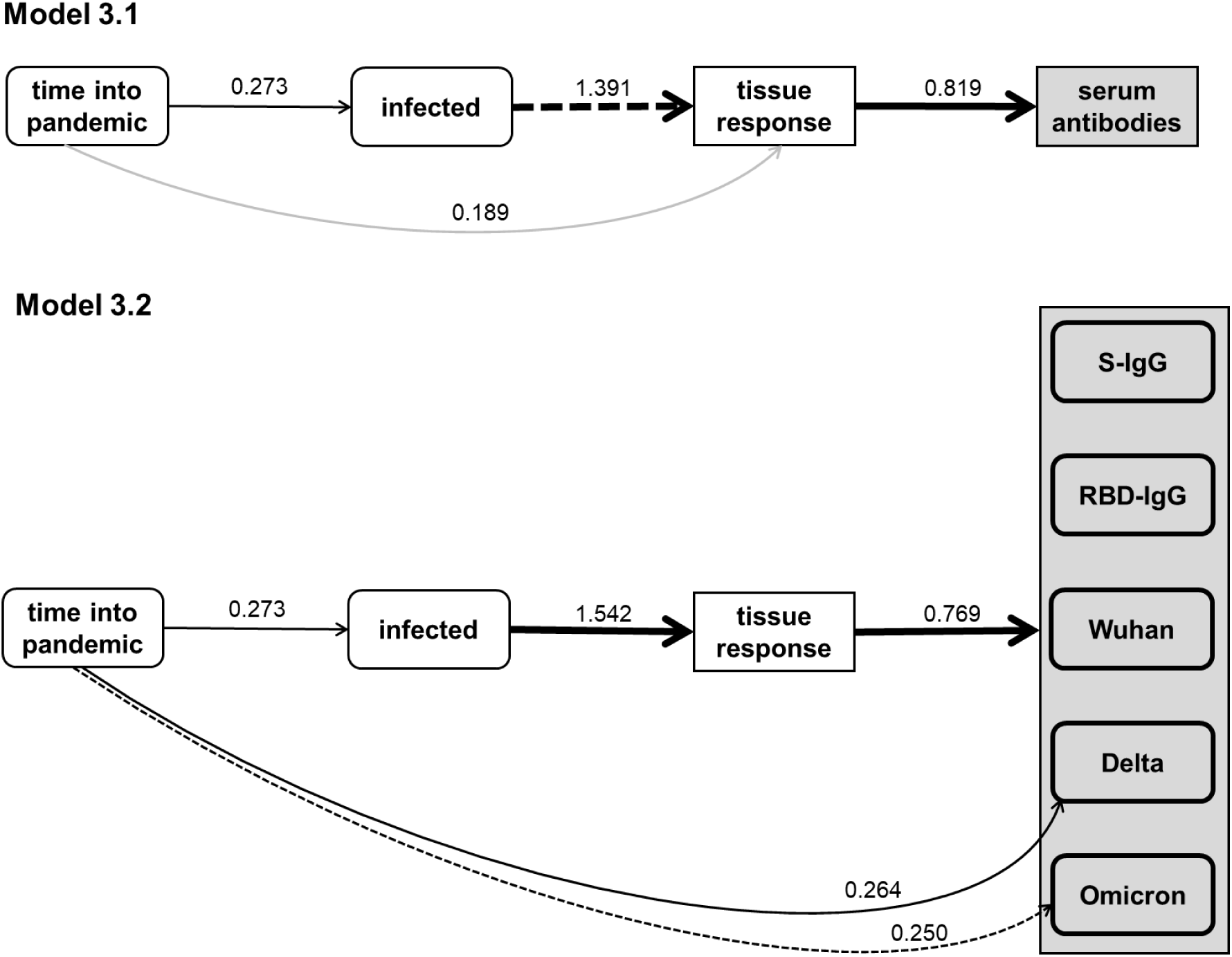
Alternative models for Model 3.

